# Long-term translocation explains population genetic structure of a recreationally fished iconic species in Japan: Combining current knowledge with reanalysis

**DOI:** 10.1101/2021.08.20.457066

**Authors:** Shuichi Kitada

## Abstract

Ayu (*Plecoglossus altivelis altivelis*) is an important freshwater fisheries resource and popular recreational fishing species in Japan, lives for only one year, and has a single breeding season. To supplement increased recreational fishing demand for this species, huge numbers of wild-born landlocked juveniles have been translocated from Lake Biwa into most Japanese rivers for more than 100 generations. Hatchery-reared juveniles born from captive-reared parents for more than 30 generations have also been extensively released. Previous studies have reported that landlocked and amphidromous forms of Ayu easily hybridise, but survival of landlocked larvae could be very low in seawater, leading to a general consensus among scientists, hatchery managers, and commercial and recreational fishers that the reproductive success of landlocked Ayu is very low, or even 0 in translocated rivers. Despite this, limited information exists regarding the reproductive success of landlocked Ayu in translocated rivers, and no study has evaluated the effects of translocation on population structure. Demonstrating that hybridisation occurs between the two forms is central to future management and conservation of this specie. To address this issue, a comprehensive literature search was undertaken, and three published genetic data sets are analysed. Analyses provide strong evidence for very high gene flow between populations, but population structure has been retained in several regions, and several populations are nested. Allele frequencies are similar in amphidromous and landlocked forms. Genetic diversity is homogeneous in amphidromous populations. Bayesian admixture analysis infers widespread introgression in Japanese rivers, with a mean introgression proportion of 24 ± 8%. Maximum likelihood admixture graphs detect two migration events from Lake Biwa to anadromous populations. Analyses consistently indicate that hybridisation between translocated landlocked juveniles and native amphidromous Ayu occurs throughout Japanese rivers.

## 1 INTRODUCTION

Anthropogenic forces often break reproductive barriers, providing an opportunity for hybridisation between closely related species that might not otherwise exist under natural conditions (Anderson, 1948, 1949; Barton, 2001; Grabenstein & Taylor, 2018; Hewitt, 1988; Hirashiki et al., 2021). A geographic area where natural hybridisation occurs is referred to as a hybrid zone, while an area where two divergent populations or closely related species occur and potentially hybridise is called a contact zone (Barton & Hewitt, 1985; Hewitt, 1988; Johannesson et al., 2020). Hybrid zones work as selective filters, enabling determination of the evolutionary roles of reproductive barriers at various stages of speciation (Barton & Hewitt, 1985; Hewitt, 1988; Martinsen et al., 2001). Repeat backcrossing and recombination of genotypes in hybrid zones results in genetic changes beyond the first hybrid generation, allowing neutral and adaptive alleles to spread among populations (Barton, 1979; Barton & Hewitt, 1985; Hewitt, 1988; Johannesson et al., 2020).

Translocation of organisms across geographic boundaries has been often used in conservation and wildlife management. It removes reproductive barriers and can lead to hybridisation of introduced and resident populations (Berger-Tal et al., 2020; Seddon et al., 2014; Taylor et al., 2021). Translocation can disturb the ecological and evolutionary processes of a species (Conant, 1988; Moritz, 1995), and unintended, negative outcomes of hybridisation can occur because of altered offspring fitness (Bucharova, 2017; Ricciardi & Simberloff, 2009). In fish management and conservation, translocations have been widely used (Allendorf et al., 2001; Bay et al., 2019; Hayes & Banish, 2017), but the consequences of this action are often poorly documented (Healy et al., 2020). Quantitative evaluation of empirical cases is needed to determine their conservation efficacy and future management efforts (George et al., 2009; Healy et al., 2020; Minckley, 1995).

Herein, a case study on Ayu (*Plecoglossus altivelis altivelis*), an iconic fish species targeted by recreational fishers in Japanese rivers, is presented. Ayu—an essentially amphidromous species, except for some land-locked populations (landlocked form) (Nishida, 1985)—can be found throughout Japan, and is one of the most well-studied fish species in the country. Ayu live for one year, and have a single breeding season (Nishida, 1986). Landlocked Ayu have been extensively translocated from Lake Biwa—the largest of landlocked populations—into amphidromous populations in rivers for more than 100 years (generations) throughout Japan. In addition, hatchery-reared amphidromous juveniles born from captively reared parents have been extensively released for more than 30 generations. Below I briefly review knowledge regarding this species and describe the aim of this study.

### 1.1 Distribution and life history of Ayu

Ayu occur widely in coastal areas of China, Korea, Japan, and Taiwan (Kwan et al., 2012), and in Vietnam (Tran et al., 2017). In China, *P. altivelis chinensis* is distributed (Xiujuan, Yunfei, & Bin, 2005). The Japanese archipelago separated from the continent of east Asia by about 17 myr ago and an intermittent gene exchange of freshwater fish species has occurred through a narrow land bridge created by lowered sea level (Taniguchi et al., 2021). Population expansion of Ayu about 130 to 390 kyr before present during the late Pleistocene shaped population structure on the continent of east Asia and the Japanese archipelago (Kwan et al., 2012). The landlocked form of Ayu in Lake Biwa had diverged from the origin of the landlocked population about 100 kyr before present (Nishida, 1985). The southernmost population of Ayu occurs on Yakushima Island, Kagoshima Prefecture (Yagishita & Kume, 2021). On Ryukyu Island, Japan, a subspecies Ryukyu Ayu (*P. a. ryukyuensis*) occurs (Nishida, 1988). Field surveys in 1986 (Nishida et al., 1992) reported wild populations of Ryukyu Ayu to occur only on Amami Oshima Island, Kagoshima Prefecture, where no population of typical Ayu was observed. Ryukyu Ayu is currently listed as an endangered species in Japan (Yagishita & Kume, 2021).

The amphidromous form of Ayu spawns in the lower reaches of rivers during autumn. Hatched larvae are immediately, passively transported downstream in currents into brackish waters, then enter the sea where they overwinter. From spring to early summer, juveniles migrate upstream. By autumn they have matured and again descend downstream to spawn, then die (Nishida, 1986). Landlocked Ayu in Lake Biwa spawn in autumn in the lower reaches of rivers that flow into the lake, after which larvae are passively transported downstream into the lake where they overwinter. This form is differentiated from amphidromous populations in ecology, morphology, and molecular characteristics (e.g., Kawanabe, 1958; Nishida, 1985, 1986; Tsukamoto & Kajihara, 1987). The spawning season of the landlocked form is earlier than that of amphidromous form (Seki & Taniguchi, 1988). In areas of central Honshu, the peak spawning season for the landlocked form is in September and that for the amphidromous form is in October (Tabata & Azuma, 1986).

### 1.2 Fisheries, translocation and hatchery releases of Ayu

Ayu is an important fisheries resource in Japanese freshwaters, and a most popular recreational fishing species. Government statistics (e-Stat, https://www.e-stat.go.jp/) indicate that Ayu is caught in almost all prefectures in Japan, with characteristically high catches along the central and southern Pacific coast (Figure S1a). The commercial catch increased markedly and peaked in 1991 before declining (Figure S1b). While statistics exclude recreational catch, this is likely to be substantial, and estimated to be ~ 5.3 × 10^6^ fish (254 tonnes) over 485 × 10^3^ days (day × fishers) in Nakagawa River, Tochigi Prefecture, alone (Kitada & Tezuka, 2002)—a tonnage similar to the commercial catch in the same year (264 tonnes) in one of the top production areas in Japan (Figure S1a). The fishing season for Ayu in this river begins on June 1 and closes at the end of October, which is a typical recreational fishing period of Ayu in Japan.

Translocation of wild-born landlocked Ayu juveniles from Lake Biwa into Tama River, Tokyo, began in 1912, and this process rapidly expanded throughout Japan in the 1950s to supplement increased recreational fisher demand (Takamura, 2009). In 1972, about 133 t (26 million juveniles, assuming ~5 g average weight (Imura, 2008)) of landlocked juveniles from Lake Biwa, Shiga Prefecture were transplanted into most rivers in which Ayu is fished in 40 of 46 prefectures (excluding Shiga Pref.) in Japan. In 1992, 524 t (105 million juveniles) of landlocked juveniles were transplanted into rivers in 41 prefectures for release (Imura, 2008). Thus, each year, huge number of juveniles of wild-born landlocked Ayu have been translocated into most prefectures and released into rivers, and this practice has occurred for more than 100 years (100 generations). The catch of wild-born landlocked juveniles in Lake Biwa increased markedly and peaked in 1991, but this also declined thereafter (Figure S1b). In the 1990s, many local fisheries cooperatives gradually changed to using hatchery-born amphidromous juveniles instead of relying on landlocked juveniles for two main reasons: an outbreak of cold-water disease in Lake Biwa (Izumi & Wakabayashi, 1997), and because landlocked forms translocated into rivers were not expected to reproduce successfully (e.g., Pastene et al., 1991; Seki and Taniguchi, 1998; Tago, 2004; Takeshima et al., 2009).

In recent years, three types of juveniles have been released: hatchery-born amphidromous fish (hatchery fish), which have been released in the largest quantity, followed by wild-born landlocked juveniles from Lake Biwa, and wild-born amphidromous fish collected in coastal waters (Figure S2a). The total number of released juveniles in Japan was about 71 million in 2020. Most wild-born juveniles are caught in Shiga Prefecture (the landlocked Ayu collected in Lake Biwa, Figure S2b), and most hatchery fish are produced in Tochigi Prefecture, followed by Miyazaki, Tokushima, Yamaguchi, and Wakayama prefectures. In hatcheries, parent fish have been captive-reared for more than 30 generations, mainly because of difficulties experienced collecting sufficient mature wild fish for synchronous spawning each year (Iguchi & Mogi, 2007). Consequently, genetic diversity of hatchery-born juveniles has decreased over time (4–31 generations) (Iguchi et al., 1999; Ikeda et al., 2005; Tsuboi et al., 2019). Every year, hatchery fish have been translocated into non-natal rivers.

### 1.3 The reproductive success of translocated Ayu and hybridisation

In the early 1990s, experiments demonstrated that amphidromous and landlocked forms of Ayu interbred easily in outdoor ponds, leading to concerns that the fitness of hybrid progeny would be reduced in the wild (Iguchi & Ito, 1994). Hybridisation in transplanted populations has remained the most serious concern for management and conservation of this species (e.g., Iguchi et al., 1997; Iwata et al., 2007; Otake et al., 2002; Seki et al., 1994; Taniguchi et al., 1983). Strontium/calcium ratios of otoliths suggest that spawning of translocated Ayu from Lake Biwa represents about 20% of the total spawning of Ayu in natural spawning grounds, with a 35% hybridisation probability (Otake et al., 2002). Genetic stock identification using allozyme markers has revealed translocated landlocked Ayu have mixed in populations, with a high proportion encountered during the high fishing season in summer (Kaewsangk et al., 2000; Seki et al., 1994; Taniguchi et al., 2002), but a lower proportion in the spawning season in autumn. Studies using allozyme and mtDNA markers have concluded that released landlocked Ayu have little effect on the genetic characteristics of native populations (Pastene et al., 1991; Seki & Taniguchi, 1998).

The mixing proportion (~25%) of the landlocked form in drifting larvae sampled in September in Sho River, Toyama Prefecture, was first estimated using a single polymorphic site of the mtDNA control region as the gene marker (Iwata et al., 2007). Mixing proportions in drifting larvae sampled in October/December and marine juveniles sampled in February/April were also estimated; the proportion of larvae of the landlocked form was < ~5% and/or close to 0% (with 95% confidence intervals including 0), respectively, indicating that Ayu juveniles translocated from Lake Biwa reproduced in rivers into which they were stocked, but that the progeny survival rate was extremely low or 0 in coastal water.

Previous results led to a consensus among Japanese scientists, hatchery managers, administrators, and commercial and recreational fishers that the reproductive success of landlocked Ayu was very low, or even 0 in translocated rivers.

Translocations and hatchery releases of Ayu have been regarded as put-and-take fisheries (e.g., Pawson, 1982; Ross & Loomis, 2001; Lorenzen et al., 2012; Beard & Mann, 2020), which provide intra-generational benefits to Ayu commercial and recreational fisheries (Tsuboi et al., 2019). Contrarily, inter-generational effects of translocations and hatchery releases and its risk assessment have received little attention for this species, despite genetic effects of hatchery-reared fish on wild populations have been a major concern, particularly in salmonid species (e.g., Araki & Schmid, 2010; Christie et al., 2014; Glover et al., 2017; Grant et al., 2017; Kitada, 2018; Laikre et al., 2010; Lorenzen et al., 2012; Ryman & Laikre, 1991; Taranger et al., 2015; Waples & Drake, 2004, and references therein). No study has evaluated the genetic effects of translocation and hatchery release on amphidromous populations within the distribution range. Demonstrating that hybridisation occurs between the two forms is central to future management and conservation of this specie.

### 1.4 Study aims

With the exception of Iwata et al. (2007), no direct evidence for the reproductive success of landlocked Ayu exists. Introgression of landlocked Ayu into amphidromous populations in Japan using neutral markers in the nuclear genome has not been identified. Here, I hypothesise that the landlocked form of Ayu hybridise with the amphidromous form in translocated rivers and cause introgression of Lake Biwa populations in Japanese rivers. To examine this hypothesis, the current status of genetic population structure is assessed, and introgression of the landlocked form into amphidromous populations is inferred throughout its Japanese distribution. Using existing (published) mtDNA and microsatellite data sets, the population structure, gene flow, genetic diversity, and admixture proportions of putative ancestry populations are analysed. Using a Bayesian clustering method on microsatellite genotypes, hybrid proportions and introgression of the landlocked form are inferred in Japanese rivers. Migration events are inferred on admxture graphs.

## 2 MATERIALS AND METHODS

### 2.1 Genetic data screening

Published genetic data were screened using the Google Scholar search system with keyword searches of “Ayu,” “genetic diversity,” “landlocked,” “*Plecoglossus*,” “population structure” and “Ryukyu Ayu.” Other known studies were also added. The population genetic structure of Ayu has been studied extensively in Japan. Initially (in the 1980s), molecular markers included allozymes (Nishida, 1985; Seki et al., 1988; Taniguchi et al., 1983), but with technological developments this expanded from the late 1990s to include mtDNA (Iguchi et al., 1997; Iguchi & Nishida, 2000; Iguchi et al., 2002; Ikeda & Taniguchi, 2002; Kwan et al., 2012; Takeshima et al., 2005; Yagishita & Kume, 2021) and microsatellite markers (Ikeda et al., 2005; Takagi et al., 1999; Takeshima et al., 2009, 2016a; Yagishita & Kume, 2021; Yamaguchi et al., 2016; Yamamoto et al., 2007) to estimate genetic characteristics of populations at regional scales. No population genetics study has used single nucleotide polymorphisms (SNPs).

Data screening identified three publicly available data sets, which covered the distribution range of Ayu. Kwan et al. (2012) inferred Ayu population structure using mtDNA control region sequences (430 base pairs) for samples from the Korean Peninsula, Japanese archipelago, and the mainland of China. Microsatellite markers have been used to describe the population structure of this species throughout its distribution based on comprehensive sampling of Japanese rivers (Takeshima et al., 2016a). Herein, the mtDNA control region *Φ*_ST_ matrix of Kwan et al. (2012) was first used to visualise population structure in east Asia. The genotypes at 12 microsatellite markers collected throughout Japan (Takeshima et al., 2016a, b) were then analysed. The overall independence of the 12 microsatellite loci was confirmed, and all loci were selectively neutral; deviations from a Hardy–Weinberg equilibrium were observed at the *Pag-070* locus, but excluding this locus did not change results (Takeshima et al., 2016a). In addition, previous data using the same 12 microsatellite markers for different samples of upstream migrants in the Yodo River system, the sole outlet to flow from Lake Biwa and the contact zone, were analysed (Takeshima et al. 2009).

### 2.2 Population structure and genetic diversity

Sample site coordinates were retrieved from the original study (Takeshima et al., 2016a), but approximate coordinates for sampling sites from Lake Biwa, Toyama Bay, and those in Kwan et al. (2012) and Takeshima et al. (2009) were obtained from Google Maps. Sample locations were plotted on a map using the ‘sf’ package in R. Genotype data were converted into Genepop format (Rousset, 2008) for implementation in the R package FinePop2_v0.2 at CRAN. Allele frequencies and expected heterozygosity (*He*) for each population were automatically estimated in the ‘read.GENEPOP’ function. Genotype frequencies were computed using Genepop on the web (Rousset, 2008). Genome-wide (averaged over loci) global *F*_ST_ values of Weir and Cockerham (1984) were computed using the ‘globalFST’ function in FinePop2_v0.2. The bias-corrected *G*ST moment estimator (Nei & Chesser, 1983, NC83) performed better than other *F*_ST_ estimators when earlier estimating pairwise *F*_ST_ in coalescent simulations (Kitada et al., 2017), so *F*_ST_ values from the NC83 estimator were computed for between pairs of populations (pairwise *F*_ST_) using the ‘pop_pairwiseFST’ function. Populations were grouped into nine geographic areas following the national fish production statistics: Hokkaido, northern Sea of Japan, western Sea of Japan, northern Pacific, central Pacific, southern Pacific, Kyushu (East China Sea), Seto Inland Sea and Lake Biwa. Pairwise *F*_ST_ were also computed for between the nine groups. An unrooted neighbour-joining (NJ) tree (Saitou & Nei, 1987) was created for the pairwise *F*_ST_ distance matrix using the ‘nj’ function in the R package ‘ape.’ The performance of the *F*_ST_ analysis applied here is evaluated by colonisation simulations and analyses of empirical data sets (Kitada et al., 2021). A network analysis using Sprittree4 v4.17.1 (Huson & Bryant, 2006) was also performed to visualise population structure.

Sampling locations were connected with pairwise *F*_ST_ values smaller than a given criterion by yellow lines to visualise gene flow. Wright’s island model assumes 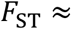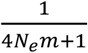, where *N_e_* is the effective population size and *m* is the average migration rate between populations (Slatkin, 1987). For example, a criterion of *F*_ST_ = 0.005 corresponds to 4*N_e_m* ≈ 199 migrants per generation (see: Waples & Gaggiotti, 2006; Whitlock & Mccauley, 1999). Isolation-by-distance relationships between populations are examined. Great circle geographic distances between all pairs of populations were computed using the ‘distm’ function in the geosphere R package, and the linear relationship between geographic and genetic distance (as measured by pairwise *F*_ST_) was estimated using the ‘lm’ function in R. Landlocked Ayu have been translocated from Lake Biwa into almost all rivers and major seed producers have sold hatchery-born juveniles to other prefectures. Waypoints (Kitada et al., 2017; Ramachandran et al., 2005), which account for natural migration paths were therefore not assumed. A Mantel correlation test was performed using the ‘mantel’ function in vegan R package with 1000 permutations.

Sample points were visualised using a colour gradient of population *He* values. The colour of population *i* was rendered as rgb (1 − *H_eO,i_*, 0, *H_eO,i_*), where *H_eO,i_* = (*H_e,i_* − min*H_e_*)/(max*H_e_* − min*H_e_*). This conversion represents the standardised magnitude of an *He* value at the sampling point, with a continuous colour gradient ranging from blue (lowest) to red (highest) for *He* values. Differences in average *He* values between all pairs of the nine geographic groups were tested by one-way ANOVA using the ‘anova’ function, with the *p*-values being adjusted for multiple comparisons with a 5% significance using the ‘TukeyHSD’ function in R.

### 2.3 Inferring admixture and hybrid individuals

Genotype data in Genepop format were converted into STRUCTURE format using PGDSpider v2.1.1.5 (Lischer & Excoffier, 2012). STRUCTURE 2.3.4 was run with no prior information about sampling locations (Pritchard et al., 2000), assuming admixture and corelated allele frequencies between populations (Falush et al., 2003). A burn-in of 100,000 sweeps followed by 500,000 MCMC sweeps was used for the number of original populations *K* = 1–5, with 10 runs for each *K*. Convergence was checked by comparing the estimated *q*-value across the 10 runs. The most likely *K* was estimated based on *ΔK* statistics (Evanno et al., 2005) using Structure Harvester (Earl & vonHoldt, 2012).

Hybridisation and hybrid individuals in contact zones can be identified by Bayesian clustering methods using highly polymorphic molecular markers such as microsatellites (Beaumont et al., 2001; Manel et al., 2005; Pierpaoli et al., 2003; Vähä & Primmer, 2006). In admixture analyses, a threshold for an individual admixture coefficient (*q*-value) inferred from STRUCTURE is generally used to identify hybrid individuals (Szatmári et al., 2021). A low threshold *q*-value of 0.1 (10%) was efficient for detecting hybrids between two distinct purebred classes or different species (Vähä & Primmer, 2006) and proved highly effective results in such empirical cases of various species (Devitt et al., 2011; Fogelqvist et al., 2015; Szatmári et al., 2021; van Wyk et al., 2017). Contrarily, the present study aims to estimate hybrids between landlocked and amphidromous forms of Ayu, which would be genetically closer; a 0.2 (20%) threshold *q*-value is used herein, where individuals with landlocked ancestral genes (0.8 > *q* > 0.2) are classified as hybrids, those with *q* ≥ 0.8 as the original landlocked form, and those for which *q* ≤ 0.2 as the original amphidromous form (Geraldes et al. 2014; Karamanlidis et al. 2021; Larison et al. 2021). The proportion of introgression of the landlocked form was estimated by summing up individuals of the original landlocked form and hybrids (*q* > 0.2) in the amphidromous populations. The threshold of *q* > 0.2 gives more conservative estimates of introgression than that of *q* > 0.1. Differences in average of introgression proportions between all pairs of the eight groups of amphidromous populations were tested using the same method as used for the ANOVA of average *He* values.

To infer migration events, we applied TreeMix (Pickrell &Pritchard, 2012) to the microsatellite data (Takeshima et al., 2016a). The analyses were performed using the nine regional groups. Using the allele frequencies and the lengths (numbers of repeats), the mean and variance in length at each locus were computed. They were used to run TreeMix v1.13. Up to five migration events were examined and the best maximum likelihood tree was found using the Akaike Information Criterion (AIC; Akaike, 1973); 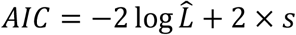, where 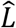 is the maximum composite likelihood and *s* is the number of parameters. In this study, *s* was three per migration event (migration edge, branch length, and migration weight). For example, for the model with two migration events, the number of parameters was six (= 2 × 3) (Kitada & Kishino, 2021).

## 3 RESULTS

### 3.1 Population structure

An unrooted NJ tree using the mtDNA control region *Φ*_ST_ matrix (Kwan et al., 2012) describes the population structure of Ayu in east Asia (Figure 1). Genetic (nucleotide) diversity (Figure 1a) was high in Japan and on the Korean Peninsula, but low in islands, Lake Biwa and in the Chinese population. Korean and Japanese populations were closely located but clearly separated each other (Figure 1b). The landlocked Ayu collected from Lake Biwa and Ryukyu Ayu collected from Amami Oshima Island had similar genetic distances from amphidromous populations. The populations on Yakushima and Amami Oshima Islands were close to the Chinese population.

**FIGURE 1.**
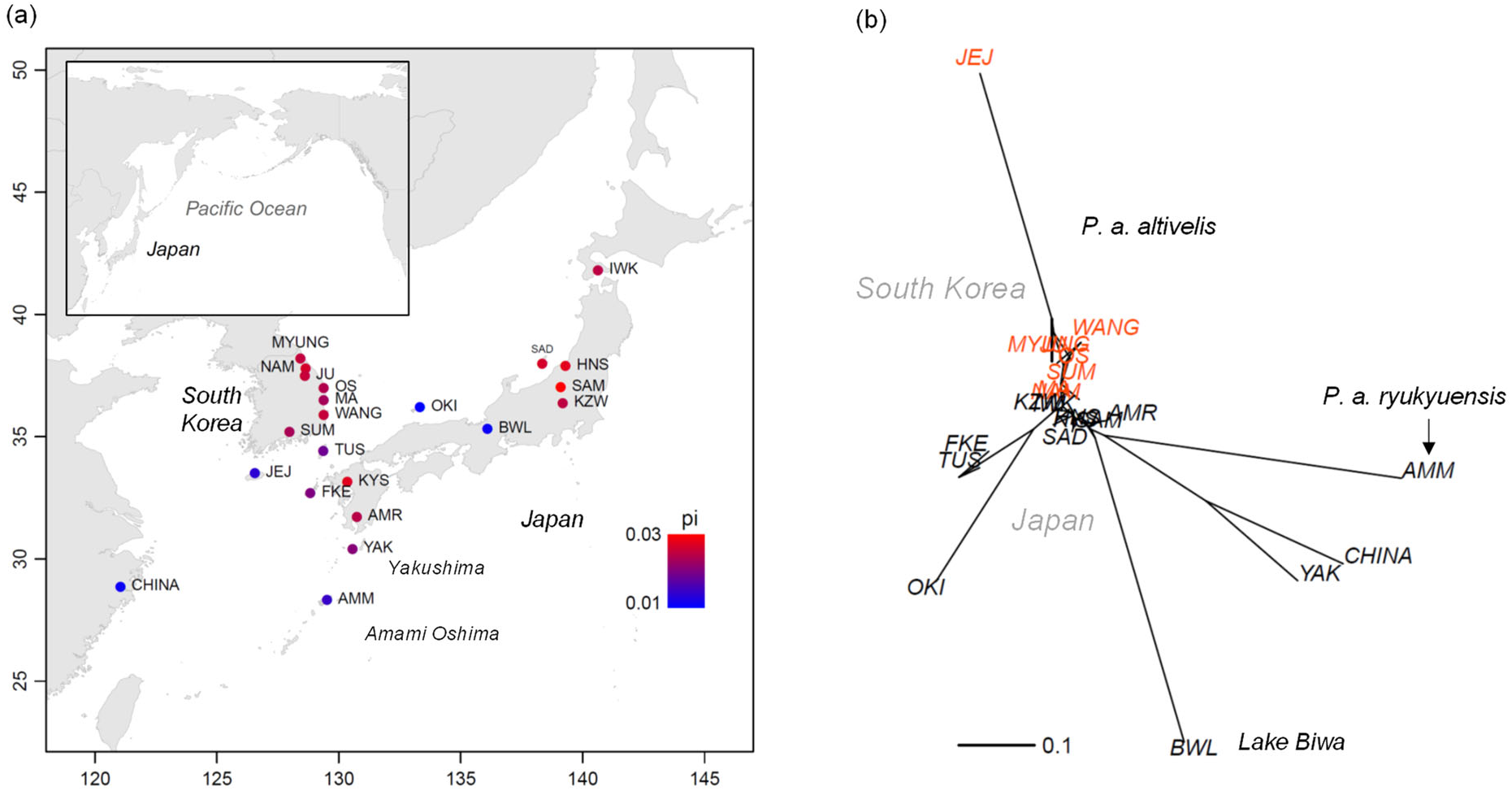
Population structure of *Plecoglossus* spp. in east Asia. (a) Sampling locations with nucleotide diversity per site. (b) Unrooted neighbour-joining (NJ) trees based on the pairwise *Φ*_ST_ matrix inferred from mtDNA control region sequences (22 populations, n = 244). Drawn from Tables 1 and 2 in Kwan et al. (2012). Drawn from Tables 1 and 2 in Kwan et al. (2012)

Genotypes of Ayu and Ryukyu Ayu at 12 microsatellite markers collected throughout Japan were analysed (120 populations, *n* = 4809, Table S1) (Takeshima et al., 2016a, b). Samples of the amphidromous form comprised upstream migrants and/or marine juveniles collected from 100 rivers/streams and or coastal waters, and individuals of the landlocked form were from 20 rivers flowing into Lake Biwa. The total sample included two Ryukyu Ayu samples collected from Amami Oshima Island. Sampling locations for the microsatellite genotyping were plotted on a map (Figure 2a). The global *F*_ST_ (± standard error, SE) was 0.0341 (± 0.0100) for all populations including Ryukyu Ayu. The mean of the pairwise *F*_ST_ was 0.0209 ± 0.0663 (standard deviation, SD) for all population pairs. The unrooted NJ tree (Figure 2b) depicted Ryukyu Ayu populations to be largely differentiated from those of Ayu, while amphidromous and Lake Biwa populations were close. The global *F*_ST_ dropped about half, 0.0162 (± 0.0074 SE) when excluding 2 Ryukyu Ayu populations. The mean of the pairwise *F*_ST_ was 0.0088 ± 0.0099 (SD) for all population pairs.

**FIGURE 2.**
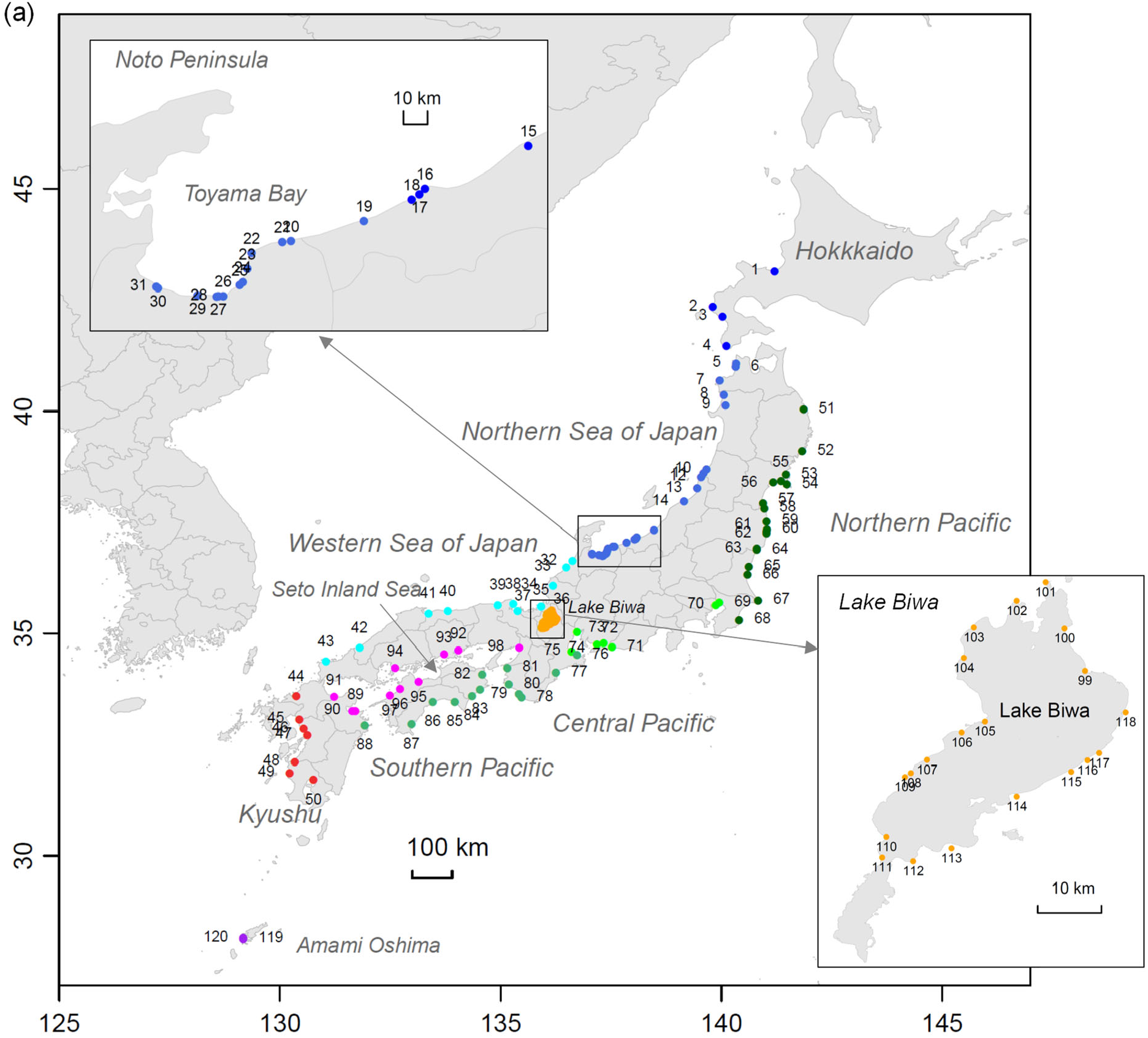
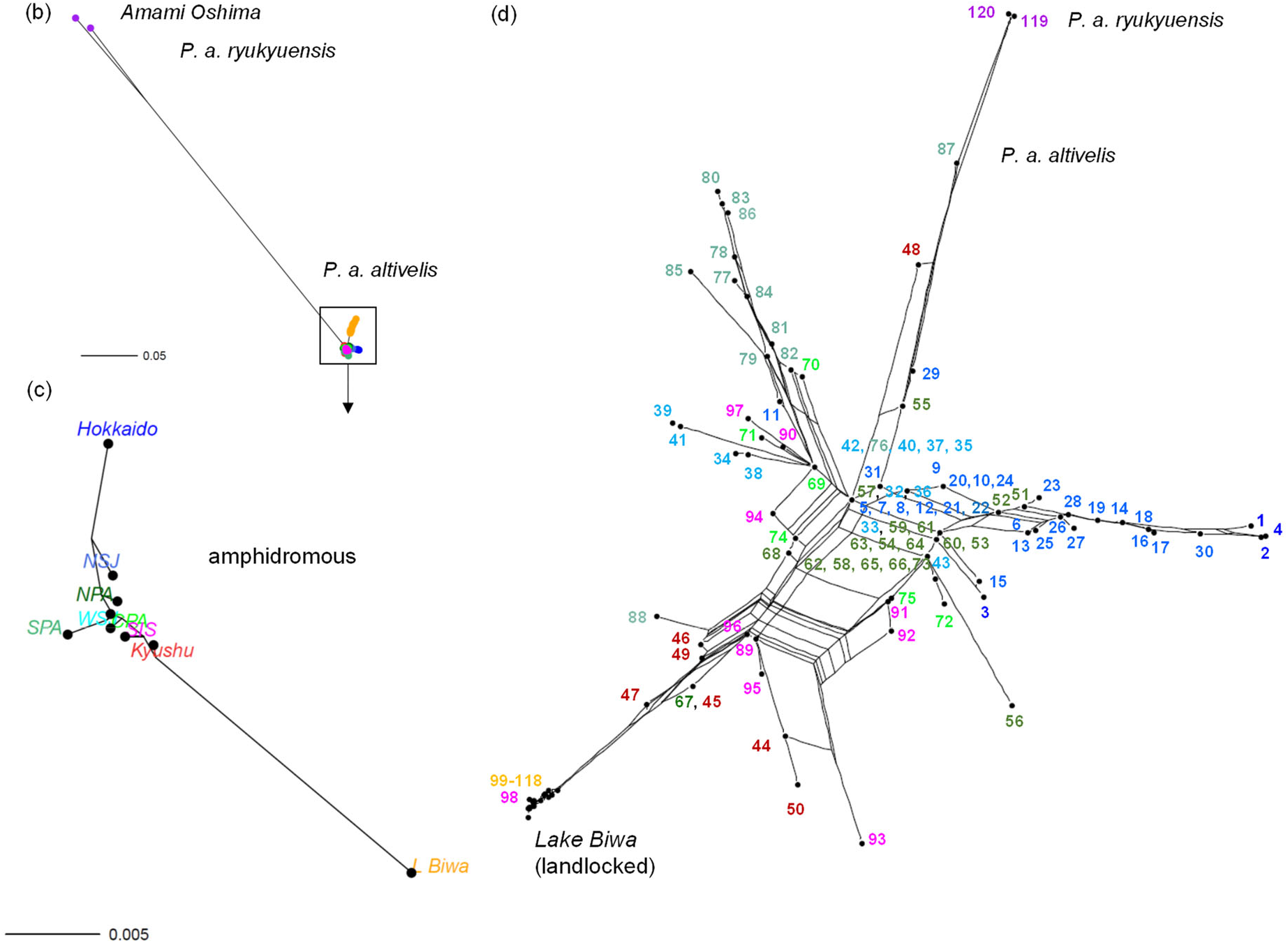
Population structure of *Plecoglossus* spp. in Japan. (a) Ayu sampling locations throughout Japan (120 populations, *n* = 4809, Table S1). Unrooted neighbour-joining (NJ) trees based on the pairwise *F*_ST_ matrix using 12 microsatellite genotypes for (b) all populations (120 populations, *n* = 4809), (c) Ayu populations grouped geographical areas. NSJ; northern Sea of Japan, WSJ; western Sea of Japan, NPA; norther Pacific, CPA; central Pacific, SPA; southern Pacific, SIS; Seto Inland Sea. (d) A neighbour network tree (Sprit tree) of 118 populations based on the pairwise *F*_ST_ matrix estimated from 12 microsatellite genotypes (n = 4746). Data are from Takeshima et al. (2016a)

An unrooted NJ tree (Figure 2b) depicted Ryukyu Ayu populations to be far from those of Ayu, while amphidromous and Lake Biwa populations were closer. Clear differences between landlocked and amphidromous forms are apparent (Figure 2c). A neighbour network tree (Figure 2d) determined reticulate divergence of amphidromous populations. Populations in Kyushu (#44–50, except 48), Midori River (#47), Ariake Sea, Kyushu and Tone River (#67), the northern Pacific were closest to landlocked populations in Lake Biwa, despite being far from Lake Biwa. Lake Biwa populations were also connected with other populations in the southern (#88) and the Seto Inland Sea (#89,93,95,96), which locate far from Lake Biwa (see Figure 2a). In other areas of the tree, reticulate networks beyond the regions were also observed. Hokkaido and the northern Sea of Japan group fall into one cluster, and the southern Pacific form a cluster. Populations from the western Sea of Japan and northern Pacific were located relatively close to each other in the centre of the tree and retained regional characteristics, with a number of nested populations. Populations from the central Pacific were nested in other clusters. The sample from Yodo River (#98), the contact zone was included in the landlocked cluster. Contrarily, the population structure (Figure S3) created using the *F*_ST_ matrix based on the same 12 microsatellite markers in Takeshima et al. (2009) located upstream migrants in Yodo River between Lake Biwa and amphidromous populations, being consistent with the original study.

Using the criterion of pairwise *F*_ST_ < 0.005 (4*N_e_m* ≈ 199), substantial gene flow between populations was apparent (Figure 3a), with very high gene flow between Ayu populations and geographic areas, but Ryukyu Ayu populations were isolated. The relationship between geographic distances and genetic distances as measured by pairwise *F*_ST_ is depicted in Figure 3b. The linear regression of *F*_ST_ on geographic distance produced an isolation-by-distance relationship within amphidromous populations (black dots). The Mantel correlation was significant (*r* = 0.32, *p* = 0.001), and *R*^2^ was 0.10. Many amphidromous populations in the within-amphidromous-population-comparison are overlapped by those in the between-Lake Biwa-comparison. Contrarily, the scatter plot (coloured dots) for Lake Biwa and amphidromous populations indicates that the linear relationship between geographic distance and genetic distance was disrupted.

**FIGURE 3.**
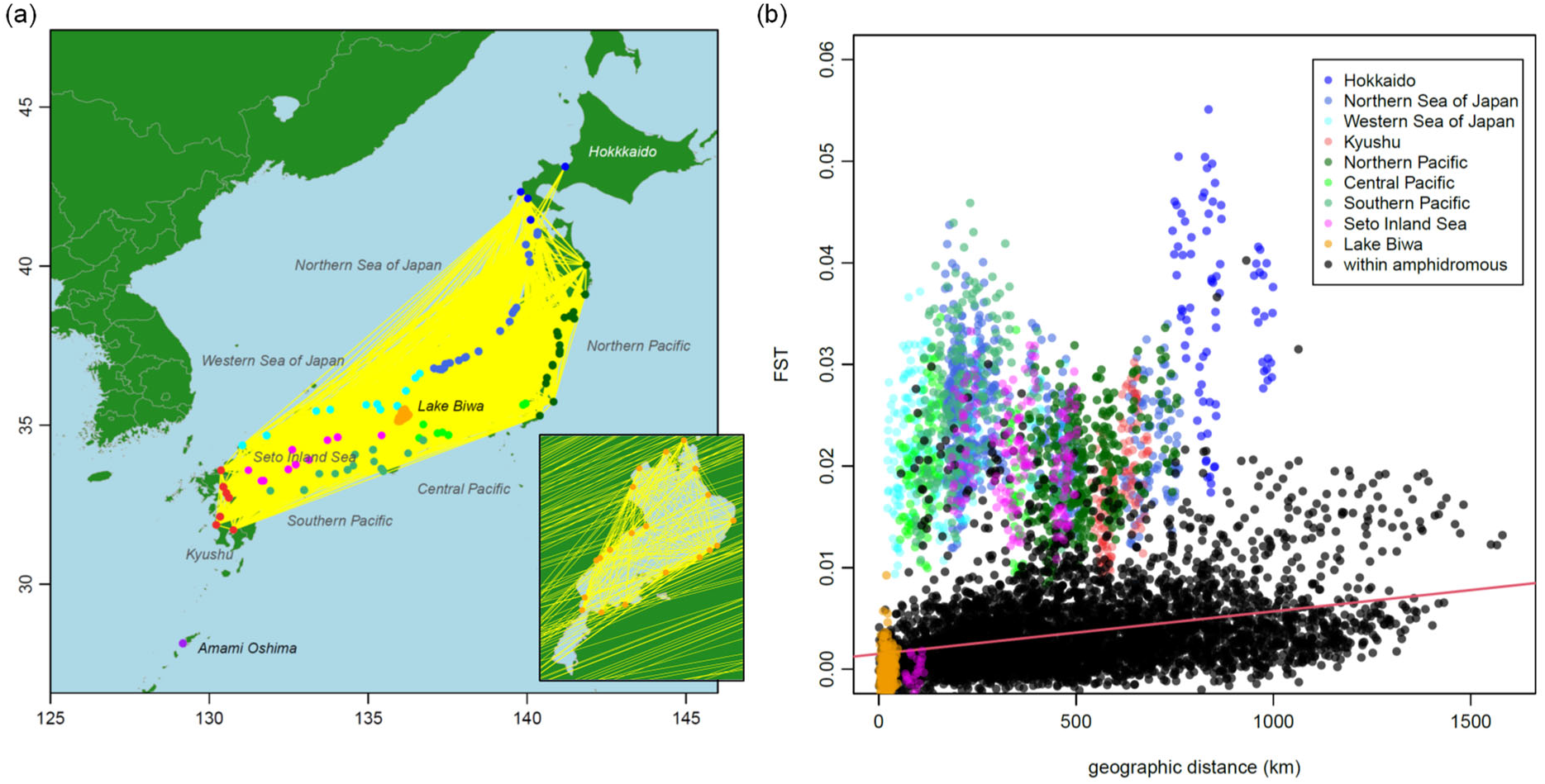
Gene flow of Ayu in Japan. (a) Map visualising geneflow between populations. Yellow lines connect population pairs with pairwise *F*_ST_ < 0.005. (b) Geographic distance vs pairwise *F*_ST_ of Ayu in Japan. Black dots depict within-amphidromous-population comparisons, with a regression line 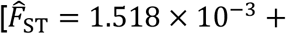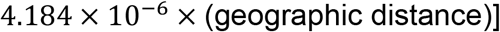 in red, with *R*^2^ 0.01; the Mantel correlation between *F*_ST_ and geographic distance is 0.32 (*p* = 0.001). Orange dots depict the samples from within the Lake Biwa population; other colours are for between Lake Biwa and amphidromous population comparisons.

### 3.2 Genetic diversity

Allele frequency distributions were surprisingly similar between amphidromous and landlocked forms of Ayu (Figures S4-1, 2), but those of Ryukyu Ayu populations differed considerably (Figure S4-3). The numbers of genotypes (Table S2) were highest in the amphidromous form at all loci. The landlocked form in Lake Biwa had substantially fewer genotypes, followed by Ryukyu Ayu. All homozygote genotypes in the landlocked form were shared with the amphidromous form, although the amphidromous form had specific homozygote genotypes. It was therefore not possible to find specific heterozygote genotypes for the first and/or early generation hybrids. In total, 1465 genotypes were found in both forms, with the amphidromous form having 1383, of which 541 (39%) were shared with the landlocked form in Lake Biwa, and the landlocked form had 618, of which 541 (88%) were found in amphidromous populations (Figure S5). Contrarily, Ryukyu Ayu had single genotypes (fixed) at six loci (Table S2), and no genotype was shared with landlocked and amphidromous forms at four of 12 loci (*Pag-002* (19 genotypes), *Pag-070* (1 genotype), *Pal-005* (2 genotypes), and *Pal-006* (1 genotype)).

Ryukyu Ayu populations had the lowest *He* values (0.15 and 0.16). After excluding these, the mean ± SD of *He* values was 0.66 ± 0.013, with a very narrow range between 0.62 and 0.68 (Figure 4a). No clear pattern in geographic distribution of *He* was apparent. Excluding Lake Biwa, *He* values were homogeneous; average *He* values in the northern and western Sea of Japan, and northern and central Pacific were significantly higher than those for Lake Biwa, while those in Hokkaido, the southern Pacific, Kyushu, and the Seto Inland Sea did not differ from those in Lake Biwa (Figure 4b).

**FIGURE 4.**
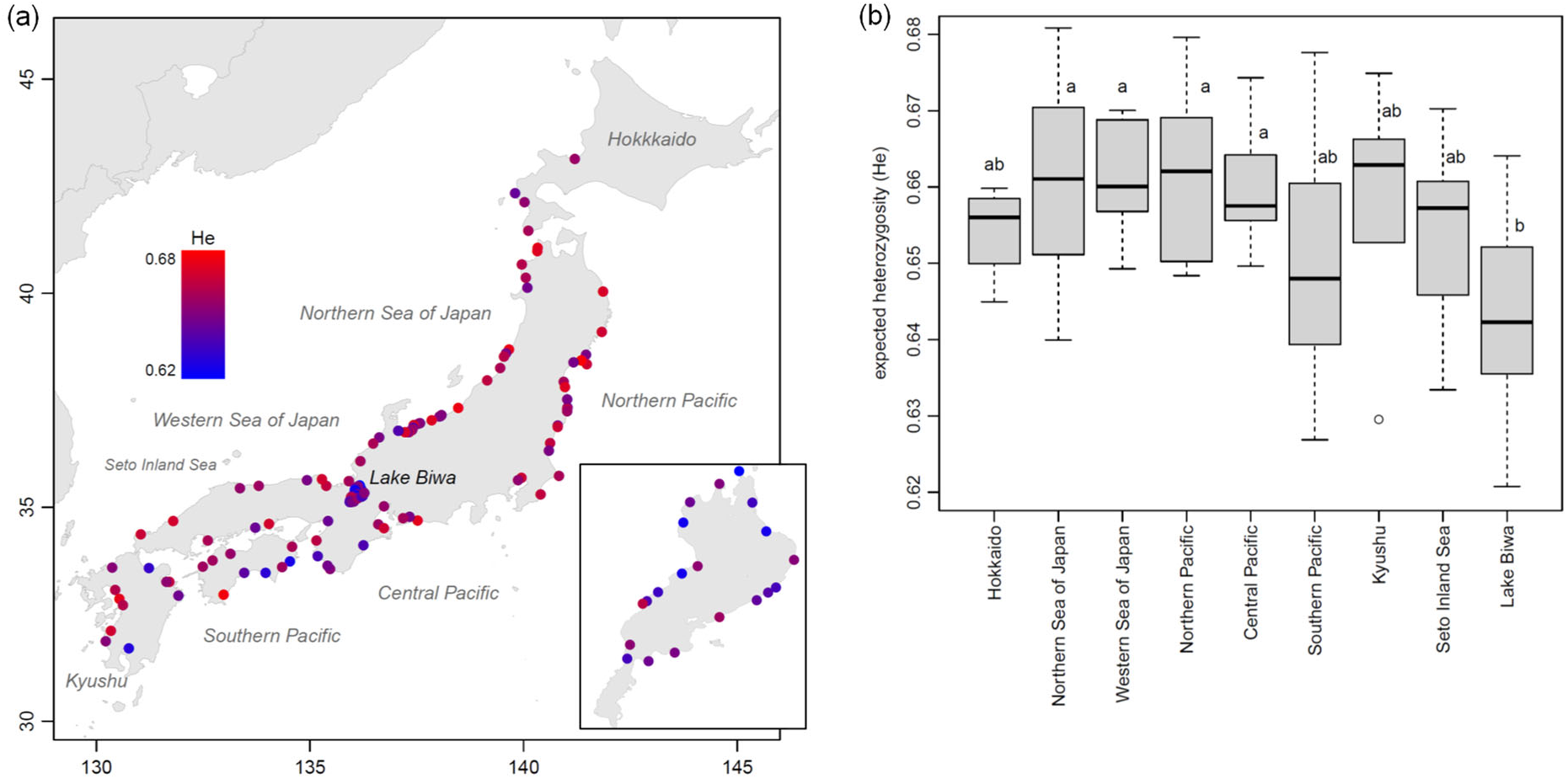
Genetic diversity of Ayu in Japan. (a) Geographic distributions of population-specific expected heterozygosity (*He*) averaged over 12 microsatellite loci in 118 populations (*n* = 4746). The colour of each population reflects the magnitude of *He* values, with a continuous colour gradient ranging from blue to red for the lowest and highest *He* values, respectively (see text). (b) Geographical distribution of *He*. Differences between areas (*p* < 0.05) are indicated by different lowercase letters. Data are from Takeshima et al. (2016a)

### 3.3 Admixture and introgression of landlocked Ayu

STRUCTURE runs suggest the most likely *K* to be 4 (Figure S6). A bar plot of individual *q*-values for *K* = 4 (Figure 5a) depicts Ayu in Japan to have four common putative ancestral origins, with similar admixture patterns within amphidromous and landlocked populations, but Ryukyu Ayu populations had different admixture pattern (purple). Blue and green colours refer to putative amphidromous ancestries, and orange to the landlocked one. The amphidromous population in Yodo River has a similar admixture pattern to that of Lake Biwa. Figure 5b depicts the geographic distribution of admixture patterns. Both forms had very similar admixture patterns within populations, but opposite distributions of *q*-values for the putative landlocked ancestry (Figure 5c, orange); the average of individual *q*-values was 78.6 ± 14.1% (SD) for Lake Biwa populations, and 21.6 ± 17.8% for amphidromous ones. Individual *q*-values of landlocked ancestry in 118 populations indicate that amphidromous and landlocked populations have similar patterns with outliers (Figure S7). For the two amphidromous ancestries (Figure 5c, green and blue), similar *q*-value distributions were obtained. The average of individual *q*-values was 39.1 ± 19.9% (median, 34.9%) for the amphidromous ancestry (green), and 39.4 ± 19.4% (median, 38.6%) for the amphidromous populations (blue). In Lake Biwa, these were 10.2 ± 7.4% and 11.2 ± 8.6%, respectively.

**FIGURE 5.**
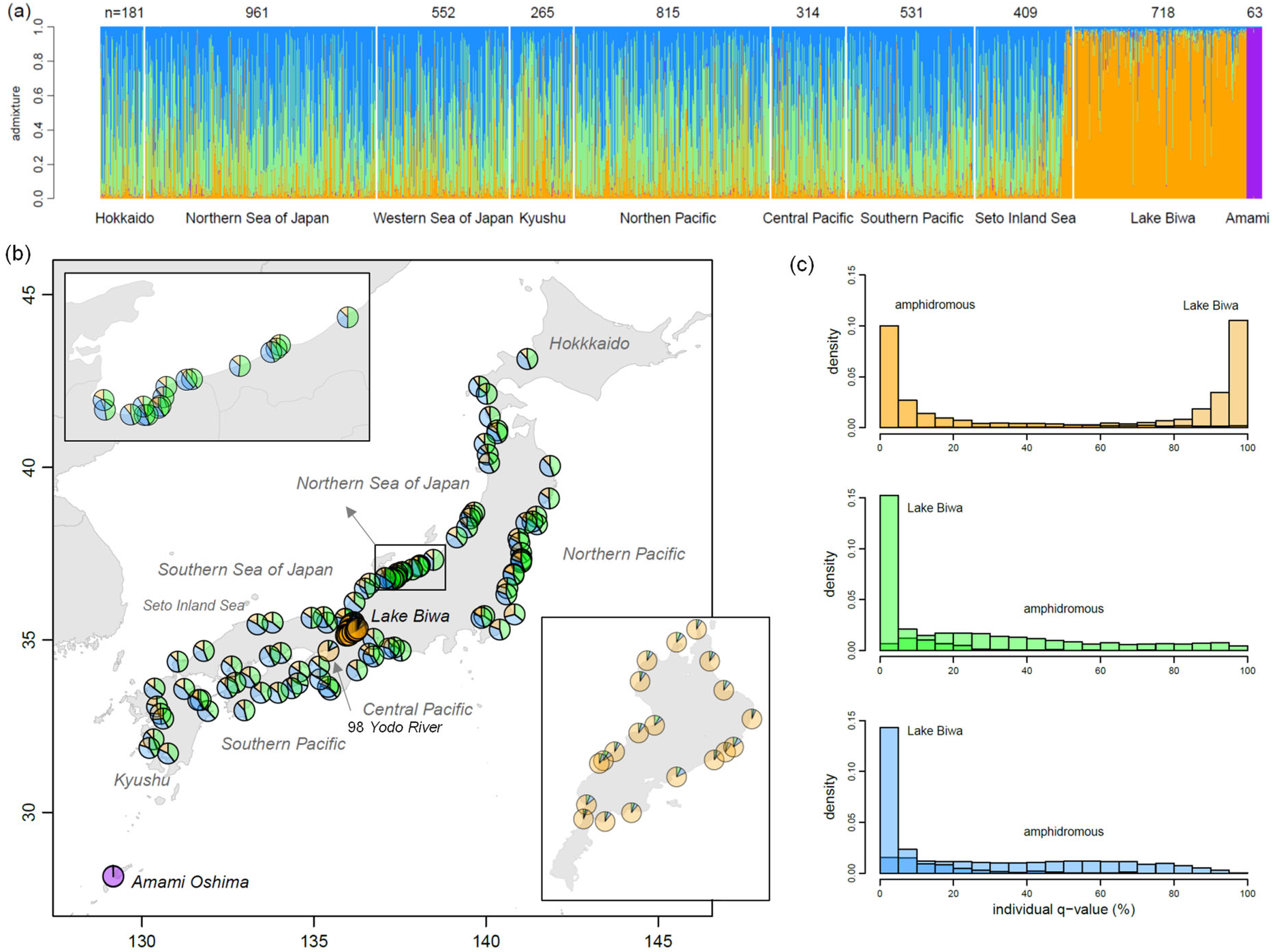
Admixture proportions in Japanese Ayu populations. (a) Coloured bars of individual *q*-values illustrate admixture proportions of inferred ancestries at *K* = 4 in 120 populations using 12 microsatellite genotypes (*n* = 4809). (b) Geographic distribution of *q*-values for the putative landlocked ancestry (orange) and the two amphidromous ancestries (green and blue) in 118 populations. (c) Distributions of the *q*-values in Lake Biwa and amphidromous populations. Data are from Takeshima et al. (2016a)

The geographic distribution of proportions of hybrid, original landlocked, and amphidromous forms using the threshold *q*-value of 0.2 (Figure 6a, Table S1) indicate hybridisation has expanded into all amphidromous populations. Average proportions were 3 ± 3% (SD) for original landlocked individuals, 21 ± 8% for hybrids, and 76 ± 8% for original amphidromous individuals in 97 amphidromous populations, excluding Yodo River, the contact zone (Figure 6b). The average proportion of introgression (original landlocked + hybrids, *q* > 0.2) was 24 ± 8% (range, 6–50%) in the amphidromous populations. A maximum of 50% were obtained in Tone River (#67) in the most southern north Pacific region, and a minimum of 6% in Sho River (#30), where Iwata et al. (2007) found that the proportion of the landlocked form was close to 0 in marine juveniles. In Yodo River (#98), the proportion of hybrids was 24%, and that of the original landlocked form 73%, with 3% original amphidromous individuals. A similar result was obtained for Lake Biwa, where the proportion of original landlocked individuals was 83 ± 7%, while that of hybrids was 15 ± 7%; similarly, 1 ± 2% amphidromous individuals were recognised.

**FIGURE 6.**
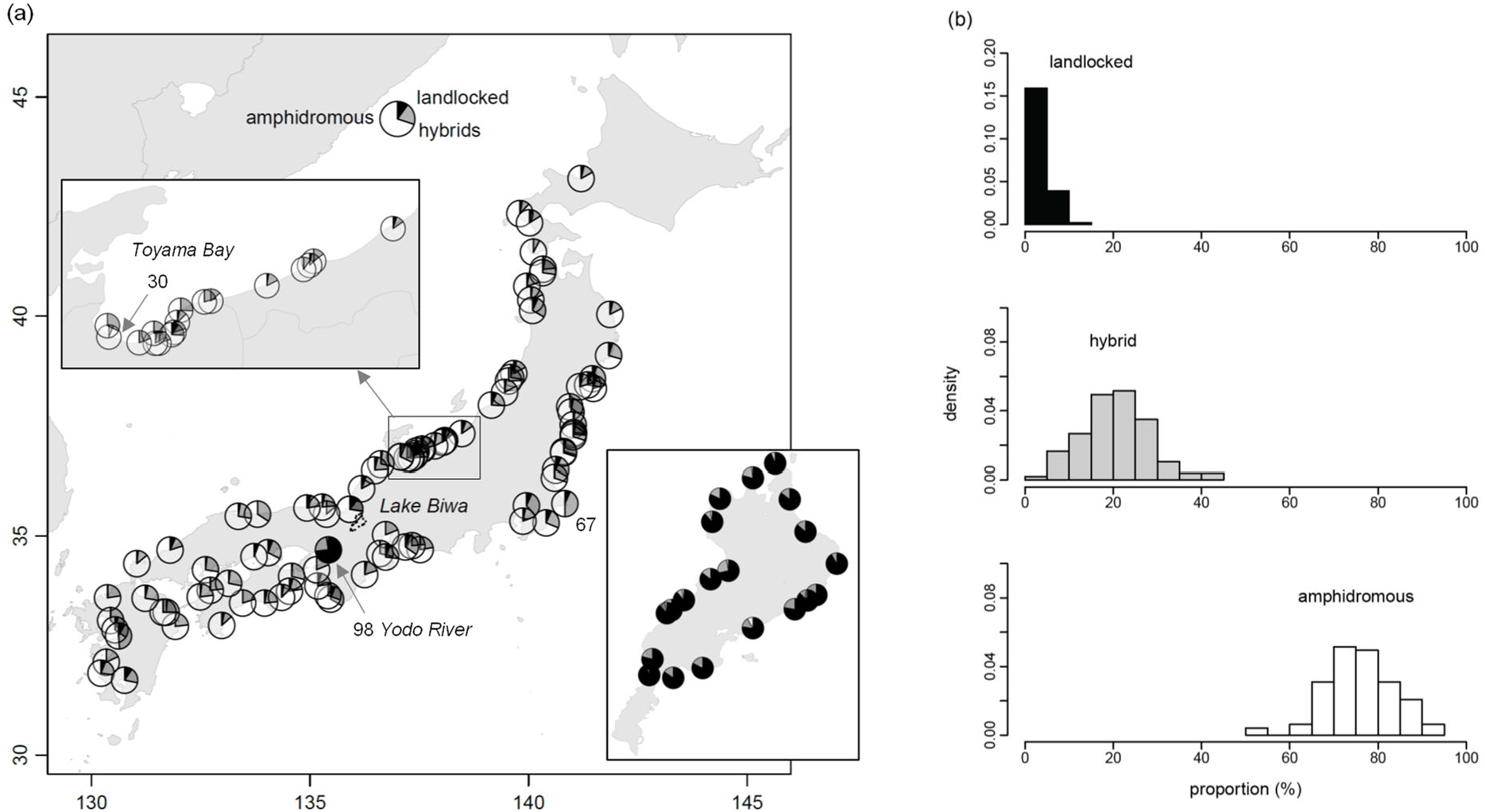
Hybrid proportions in Japanese Ayu populations. (a) Map of proportions of hybrid and original forms in 118 populations (*n* = 4746). Numbers: 30, Sho River; 67, Tone River (see text). (b) River-specific % individuals of hybrids and original forms in 97 amphidromous populations, excluding Yodo River. Data are from Takeshima et al. (2016a)

The average proportion of introgression was lower in Hokkaido, while higher in northern Pacific and Kyushu, but introgression proportions were homogeneous in all regions excluding Hokkaido and the northern Sea of Japan (Figures 7a, S8). In the TreeMix analysis, Lake Biwa was used as a root population considering the population structure (Figures 1,2). The most parsimonious maximum likelihood rooted tree (Figures 7b, S9) indicated that the divergence of Lake Biwa was similar with amphidromous populations, and two migration events, from Lake Biwa to northern Sea of Japan and Hokkaido were detected, which were consistently found for more than 20 runs.

**FIGURE 7.**
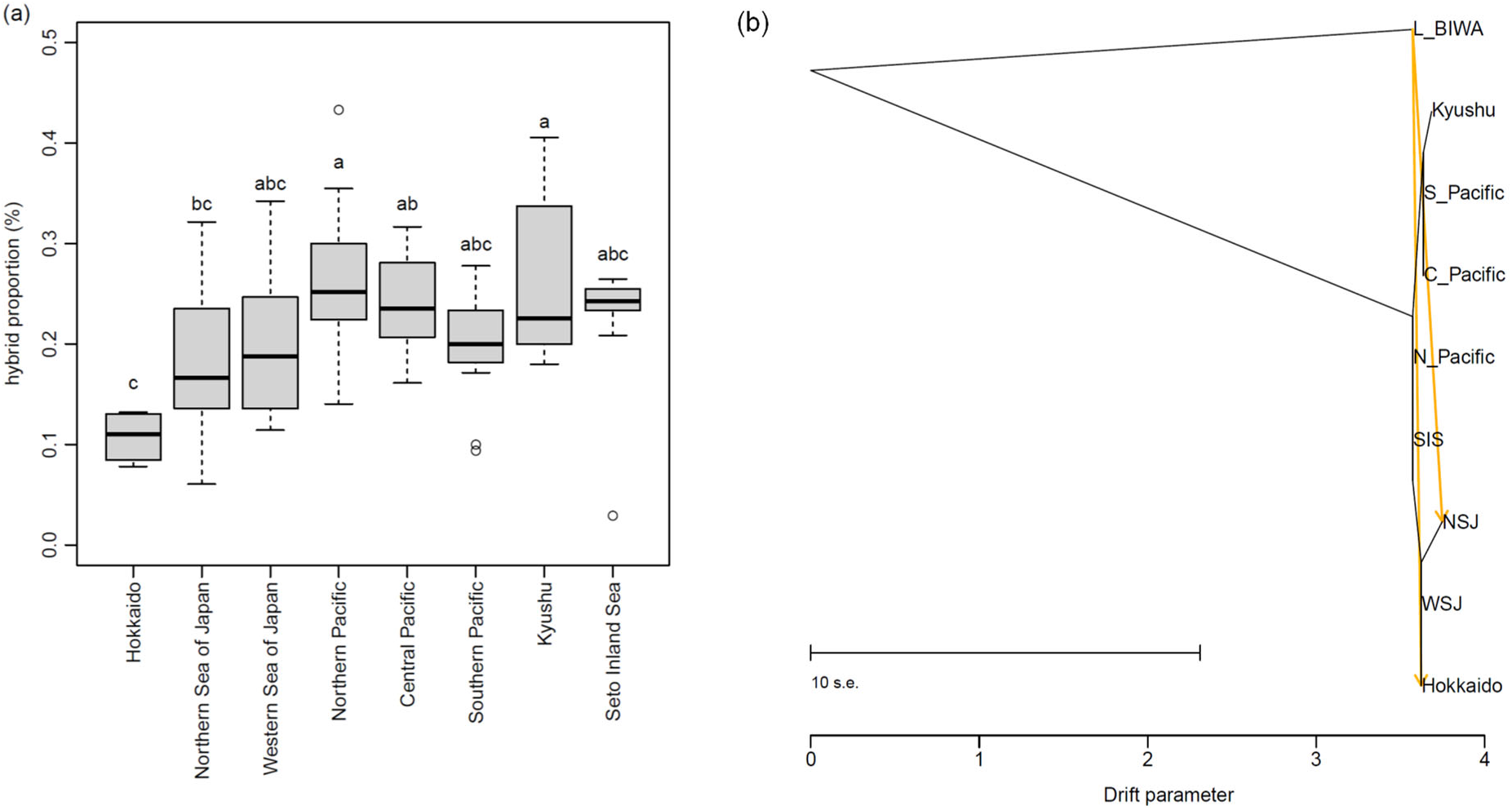
Introgression of landlocked Ayu into amphidromous populations. (a) Proportions of introgression by geographical region. Differences between areas (*p* < 0.05) are indicated by different lowercase letters. (b) The most parsimonious maximum likelihood rooted tree using TreeMix based on 12 microsatellite loci in 118 Ayu populations in Japan. NSJ; northern Sea of Japan, WSJ; western Sea of Japan, SIS; Seto Inland Sea. Data are from Takeshima et al. (2016a)

## 4 DISCUSSION

Analysed sample data were obtained from upstream migrant and/or marine juvenile Ayu (Takeshima et al., 2016a). With the exception of the sample from Yodo River, no amphidromous population was included in the cluster of Lake Biwa (Figure 2d), and the proportions of the original landlocked form were 0 in most amphidromous populations (Figure 6, Table S1). Results indicate that the probability of samples containing released landlocked juveniles is very low.

### 4.1 Population structure and translocation

The large discrepancy between population structures inferred from the mtDNA control region (Figure 1) and microsatellite markers (Figures 2) indicates that the genetic difference between amphidromous and Lake Biwa Ayu was close, when considering both maternal and paternal inheritance. The population structure (Figures 2c,d) tends to be maintained in Hokkaido, the northern Sea of Japan, Kyushu, and the southern Pacific, despite very high gene flow between populations and areas (Figures 3a).

Juvenile Ayu spend months at sea over winter (Nishida, 1986), and drift in sea currents and/or move to natal and adjacent river mouths (Iguchi et al., 2006). Juveniles commence their upstream migration in spring at ~ 60 mm body length (Tsukamoto & Kajihara, 1987); those individuals that stray, that do not return to natal rivers, contribute to gene exchange among adjacent rivers. Long-range dispersal to offshore waters might reduce survival of marine juveniles, resulting in environmental-induced natal river fidelity (Iguchi et al., 2006). This should contribute to population structuring despite high gene flow. Temporally stable and differentiated structures with high gene flow (Hauser & Carvalho, 2008) were reported for Atlantic cod (*Gadus morhua*) populations (*F*_ST_ = 0.0023; Knutsen et al., 2003), Atlantic herring (*Clupea harengus*) in Swedish waters (*F*_ST_ = 0.002–0.003; Larsson et al., 2010), and Pacific herring (*C. pallasii*) in British Columbia (*F*_ST_ = 0.003; Beacham et al., 2008). Results for Ayu population structure in some regions, despite the high gene flow (*F*_ST_ = 0.0162), are comparable to these findings for cod and herring. Contrarily, genetic differentiation was very low in the western Sea of Japan, northern and central Pacific, and Seto Inland Sea, with many populations nested among clusters (Figures 2d).

Allele frequency distributions were similar between amphidromous and landlocked Ayu forms (Figure S4), and the amphidromous form shared 541 (39% of 1383) of genotypes with the landlocked form, while 541 (88% of 618) genotypes in the landlocked form in Lake Biwa were found in amphidromous populations (Figure S5). The isolation-by-distance relationship within 98 amphidromous populations (Figure 3b) agrees with those inferred from 49 populations on the Sea of Japan coast and from 39 populations on the Pacific coast (Takeshima et al., 2016), showing the maintenance of population structure in several regions. However, the linear relationship between geographic distance and genetic distance was disrupted, with many amphidromous populations in the within-amphidromous-population-comparison overlapped by those in the between-Lake Biwa-comparison. The coefficient of variation of *He* values was very low at 2%, indicating the homogeneous distribution of *He* in Japan (Figure 4). This result similarly suggests that the allele frequencies of amphidromous populations were altered by translocation of landlocked fish.

### 4.2 Hybridisation between landlocked and amphidromous forms

The sample from Yodo River (#98), the contact zone was included in the landlocked cluster, with the proportion of hybrid and original landlocked individuals within it being very similar to those in Lake Biwa, in that there were no original amphidromous individuals. Upstream migration of Ayu along Yodo River into Lake Biwa, the sole outlet to flow from Lake Biwa, was interrupted by the construction of three dams, while well-developed larvae have been supplied from Lake Biwa into this contact zone every year, which might survive in coastal waters and return to migrate up Yodo River (Takeshima et al., 2009). Results agree with speculation by Takeshima et al. (2009), in that sample #98 collected from Yodo River was supplied from Lake Biwa and survived in coastal waters during winter. However, the unrooted NJ tree based on different samples (Takeshima et al., 2009) (Figure S3) located samples of upstream migrants between Lake Biwa and amphidromous populations, suggesting hybridisation occurred in the Yodo River system. The admixture and hybrid proportions (Figures 5, 6) in Lake Biwa should reflect the demographic history of the landlocked form, which might have been naturally invaded by amphidromous ancestry about 100 kyr before present (Nishida, 1985).

Tone River (#67), located a short distance from the amphidromous cluster in the neighbour network tree (Figure 2d) had the highest introgression proportion (50%) of all amphidromous populations (Table S1, Figure S8). Each year throughout the 1970s more than 1 × 10^6^ landlocked Ayu juveniles were translocated from Lake Biwa into a tributary of Tone River (Takamura, 2009). Natural recruitment of amphidromous Ayu depends primarily on the coastal environment, particularly that of estuaries (Takahashi et al., 1998; Tago, 2002). The substantial catch in this area (Figure S1) suggests an appropriate estuarine environment for juvenile survival, which might enhance hybrid fitness. Midori River in Ariake Sea (#47) also had the second highest introgression proportion (49%), where large estuaries exist. Contrarily, in Sho River (#30) in Toyama Bay, the introgression proportion was the lowest (6%) for upstream migrants, where Iwata et al. (2007) found that the proportion of the landlocked form was < ~5% in October/December and/or close to 0% in marine juveniles sampled in February/April. The unique environment of Toyama Bay, with a small tidal range about 20–30 cm (Tago & Watanabe, 2009) and a depth exceeding 1000 m (Watanabe, Ohtsu, & Otsuki, 2000) may have caused the very low reproductive success of landlocked Ayu in this area, with small catch (Figure S1a). Moreover, the mtDNA marker of a single-site marker in the mtDNA control region could detect a maternal but not paternal effect on reproduction of the landlocked form (Iwata et al., 2007).

Marine juveniles collected in coastal waters in late March and early April (samples #37, 38, 62) comprised 14%, 20%, and 27% hybrids, respectively (Table S1), suggesting that juvenile hybrids survived at sea. Experimental tests measured survival rates of Ayu larvae in seawater 20 days after hatching in freshwater (Figure S10a) (Tabata & Azuma, 1986). The average survival rate in 18°C (and 23°C) seawater temperature was 82% (46%) for the amphidromous form, 60% (22%) for the landlocked *strain*, and 59% (5%) for the landlocked form. In the experiment, parents collected from rivers in October which produced hatchlings of 6–7 mm total length (TL) were defined as amphidromous form, while those which collected in September and produced hatchlings of 5 mm TL were defined as the landlocked *strain*. Landlocked form larvae were born from wild parents collected in Lake Biwa. Survival rates (Figure S10a) were similar for the landlocked strain and landlocked form at 18°C, but intermediate for landlocked strain at 23°C, suggesting that landlocked strain parent fish included hybrids. Iguchi and Yamaguchi (1994) found that 24-hour survival rates of amphidromous hatchlings were 76 ± 13% in 80% artificial sea water (20°C), while those of the landlocked-form were significantly low at 34 ± 20% (Figure S10b). These results suggest that hybrids can survive in brackish and sea waters, though the survival rates are lower than for the pure amphidromous form.

The landlocked form of Ayu in Lake Biwa had diverged from the origin about 100 kyr before present (Nishida, 1985). Well-developed larvae have been supplied from Lake Biwa into Yodo River every year, the sole outlet to flow from Lake Biwa (Takeshima et al., 2009). Natural geneflow from Lake Biwa to amphidromous populations is therefore extremely limited. Contrarily, more than a million of landlocked juveniles were transplanted into rivers in most prefectures (Imura, 2008). In northern Sea of Japan, every year more than millions of juveniles had been transplanted from Lake Biwa to Toyama Bay until around 2000 (Tago, 1999, 2004). In Hokkaido, 50 thousand of landlocked Ayu juveniles were first transplanted from Lake Biwa to Yoichi River in 1964, and the success leaded to translocations to other rivers in 1965, namely Atsuta, Toshibetsu and Nikanbetsu Rivers (https://www.town.yoichi.hokkaido.jp/machi/yoichistory/2004/sono6.html). The two migration events, from Lake Biwa to northern Sea of Japan and Hokkaido amphidromous populations found in the maximum likelihood rooted tree suggest that the admixture was caused by human-induced translocations (Figures 7b, S9).

## 5 CONCLUSIONS

Analyses in this study consistently indicate that translocated landlocked juveniles have reproduced with native amphidromous Ayu in Japanese rivers and altered population structure. Widespread introgression in Japanese rivers is inferred, with mean introgression proportion of 24 ± 8%. An important action to ensure Ayu sustainability should be to avoid the use of translocated landlocked Ayu juveniles to stock rivers. Further experiments should be needed to estimate survival rates of hybrids in various coastal environments. Rivers or areas where there was no release of landlocked or hatchery-reared Ayu within a prefecture would be useful to limit gene flow and dilute any genetic effects of translocations and hatchery releases through backcrosses with wild individuals (Anderson, 1949; McGinnity et al., 2003; Hindar et al., 2006; Tymchuk et al., 2006; Roberge et al., 2008; Kitada et al., 2019). Healthy stocks of wild fish are necessary for no-release rivers and/or areas to succeed as a management option. Fortunately, wild fish genes remain dominant in Ayu populations, with 76 ± 8% of individuals being originally amphidromous. Indeed, management commenced when the Shubutogawa Fishers Cooperatives, Hokkaido, stopped the translocation of Ayu juveniles from Honshu (which began in 1984) in 2013 to avoid genetic risks to the conservation of this species at its northern margin. Efforts to create and/or restore spawning grounds in this river have continued in collaboration with fishers and locals (https://www.shubutogawa.com/), indicating future direction of management and conservation of this species.

## Supporting information

Supplemental Information, and will be used for the link to the file on the preprint site.

## Acknowledgements

This research was made possible by access to microsatellite genotype data for Ayu maintained by Hirohiko Takeshima and his colleagues. Gratitude is also expressed to the researchers and staff responsible for the time-consuming sampling of Ayu throughout its distribution in Japan that enabled this dataset to be generated. This study was supported by the Japan Society for the Promotion of Science Grant-in-Aid for Scientific Research (KAKENHI) No. 18K0578116. Steve O’Shea, PhD, from Edanz is also thanked for editing a draft of this manuscript.

## SUPPORTING INFORMATION

Additional supporting information may be found online in the Supporting Information section.

## DATA AVAILABILITY STATEMENT

All source data used are in the public sector, and links to their online sources are specified in the text or in Supporting information.

