## Supplemental Information, and will be used for the link to the file on the preprint site. for "Long-term translocation explains population genetic structure of a recreationally fished iconic species in Japan: Combining current knowledge with reanalysis"

Shuichi Kitada

This PDF file includes
Supplemental Figures S1–S10.
Supplemental Tables S1 and S2 are presented as an Excel file.

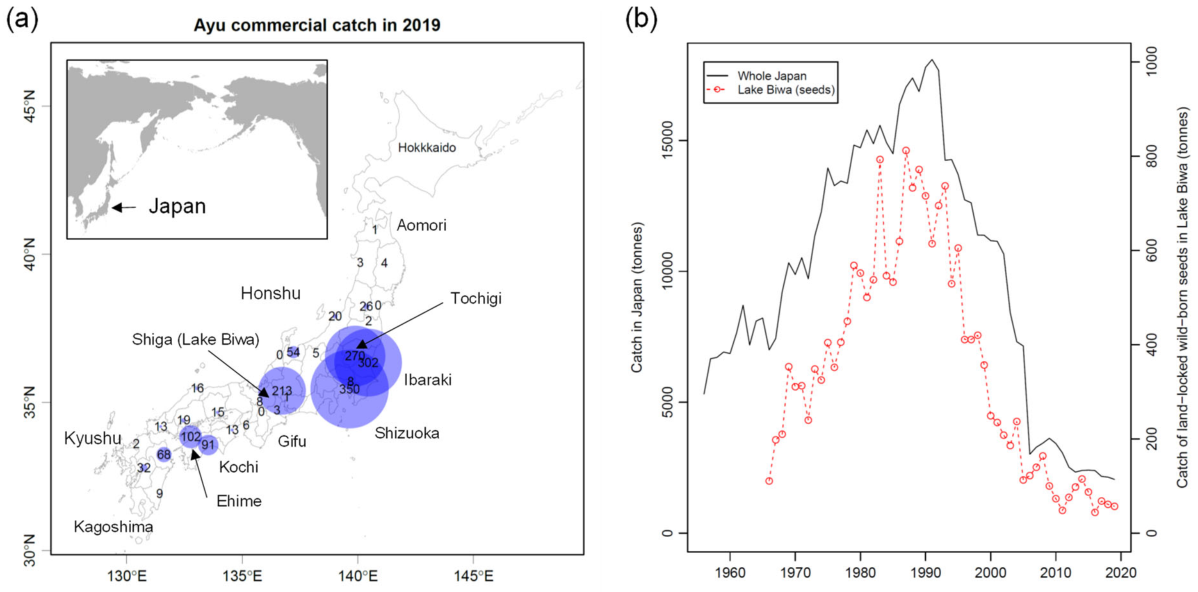


**Figure S1** Ayu catch and release statistics in Japan. (a) Commercial catch by prefecture (metric tonnes). Circle sizes are proportional to catch. (b) Commercial catch throughout Japan, and catch of wild-born juveniles in Lake Biwa, Shiga Prefecture. Retrieved from e-Stat Statistics of Japan, https://www.e-stat.go.jp/ accessed on July 27, 2021, and from Shiga Prefectural Government statistics, https://www.pref.shiga.lg.jp/, accessed on July 1, 2021

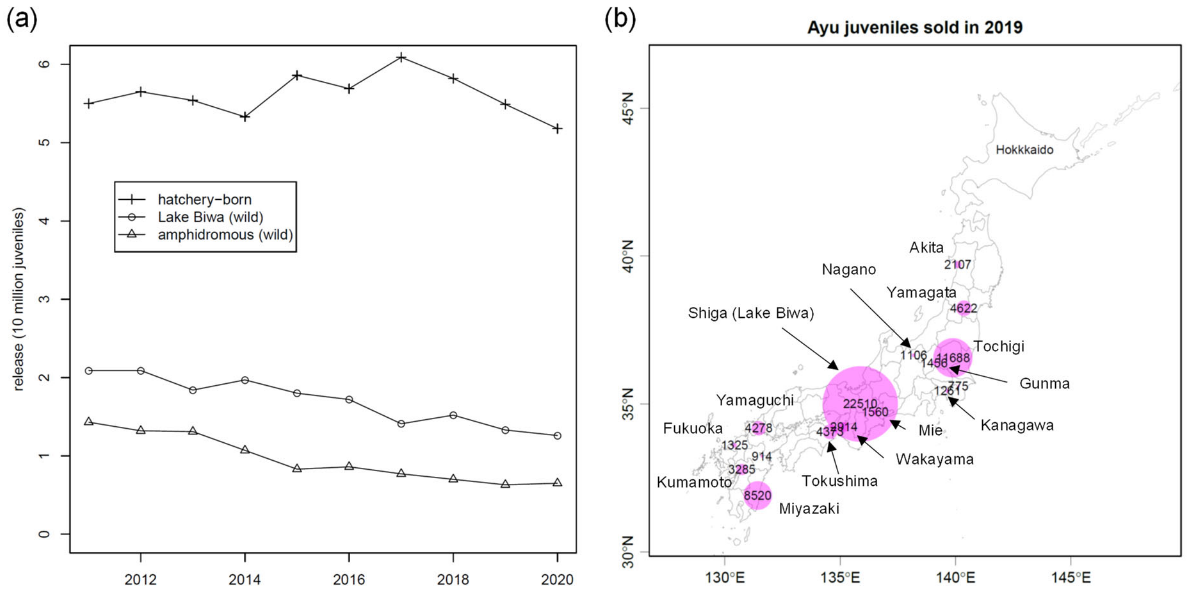


**Figure S2** Juvenile release and production statistics. (a) Changes in the number of released seeds. Estimated using the release statistics in weight from National Federation of Inlandwater Fisheries Cooperatives (http://www.naisuimen.or.jp/ accessed on July 12, 2021), assuming 10 g per juveniles (https://www.jfa.maff.go.jp/j/enoki/attach/pdf/naisuimeninfo-11.pdf). (b) Juveniles sold by prefecture (× $10^3$ fish, retrieved from e-Stat Statistics of Japan, https://www.e-stat.go.jp/ accessed on July 27, 2021)

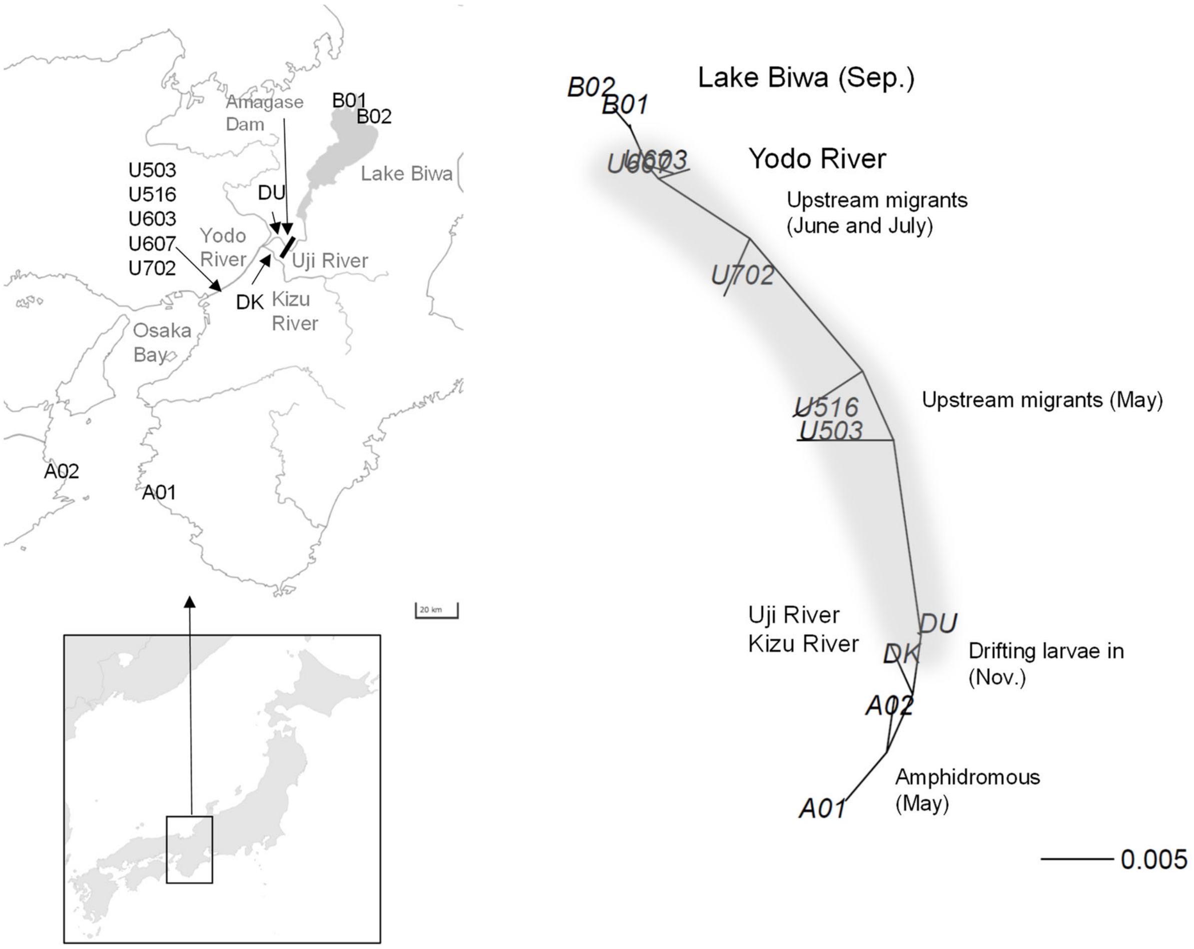


**Figure S3** Unrooted neighbour-joining (NJ) trees for Ayu in the Yodo River system. Drawn from the pairwise $F_{ST}$ matrix in Table 3 in Takeshima et al. (2009). Upstream migrants (U503, U516, U702) were located between Lake Biwa and amphidromous populations, consistent with the original study

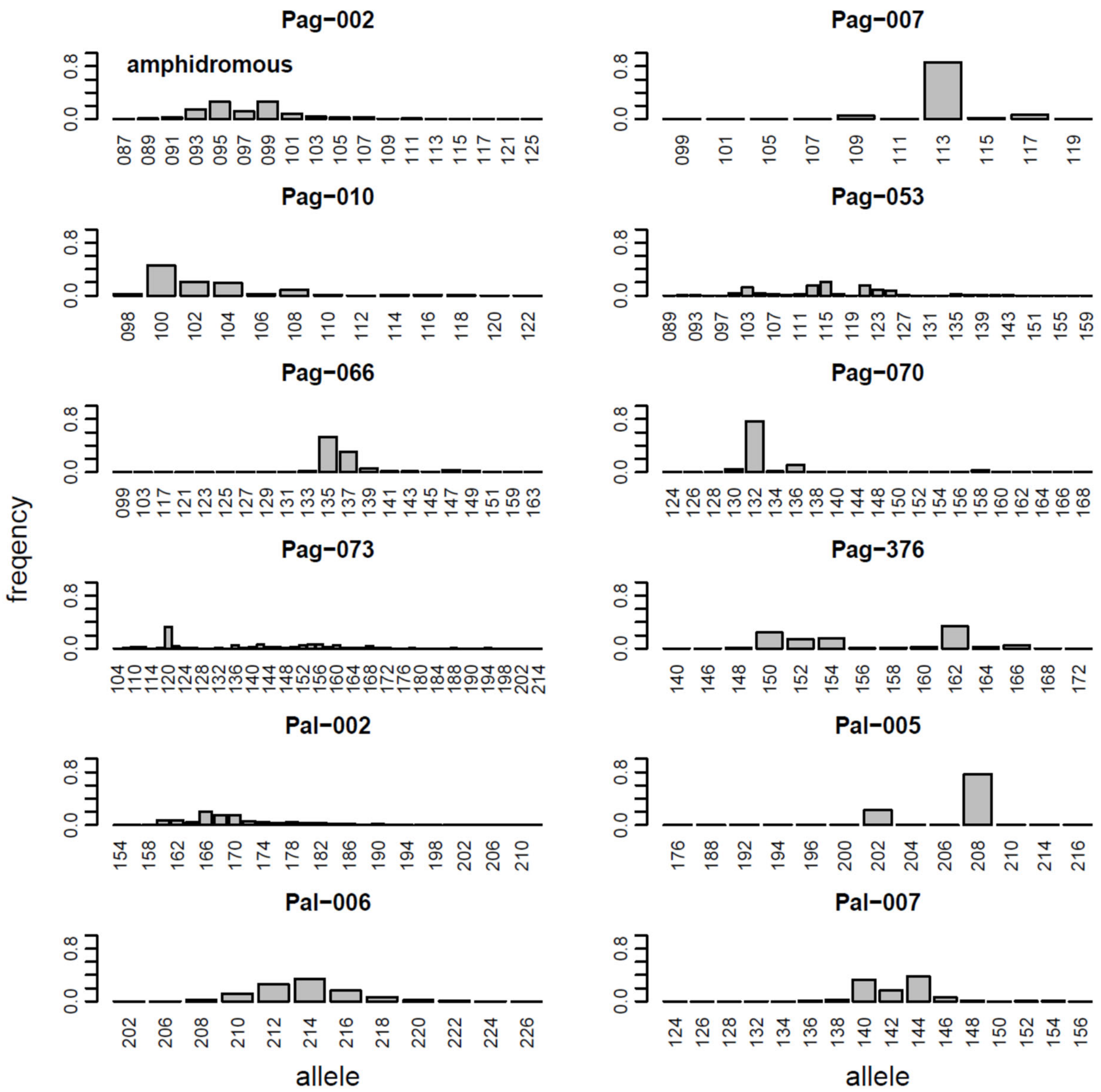


**Figure S4-1** Mean allele frequencies of amphidromous form of Ayu at 12 microsatellite loci (97 populations, $n$ = 3987). Yodo River is excluded. Data are from Takeshima et al. (2016a)

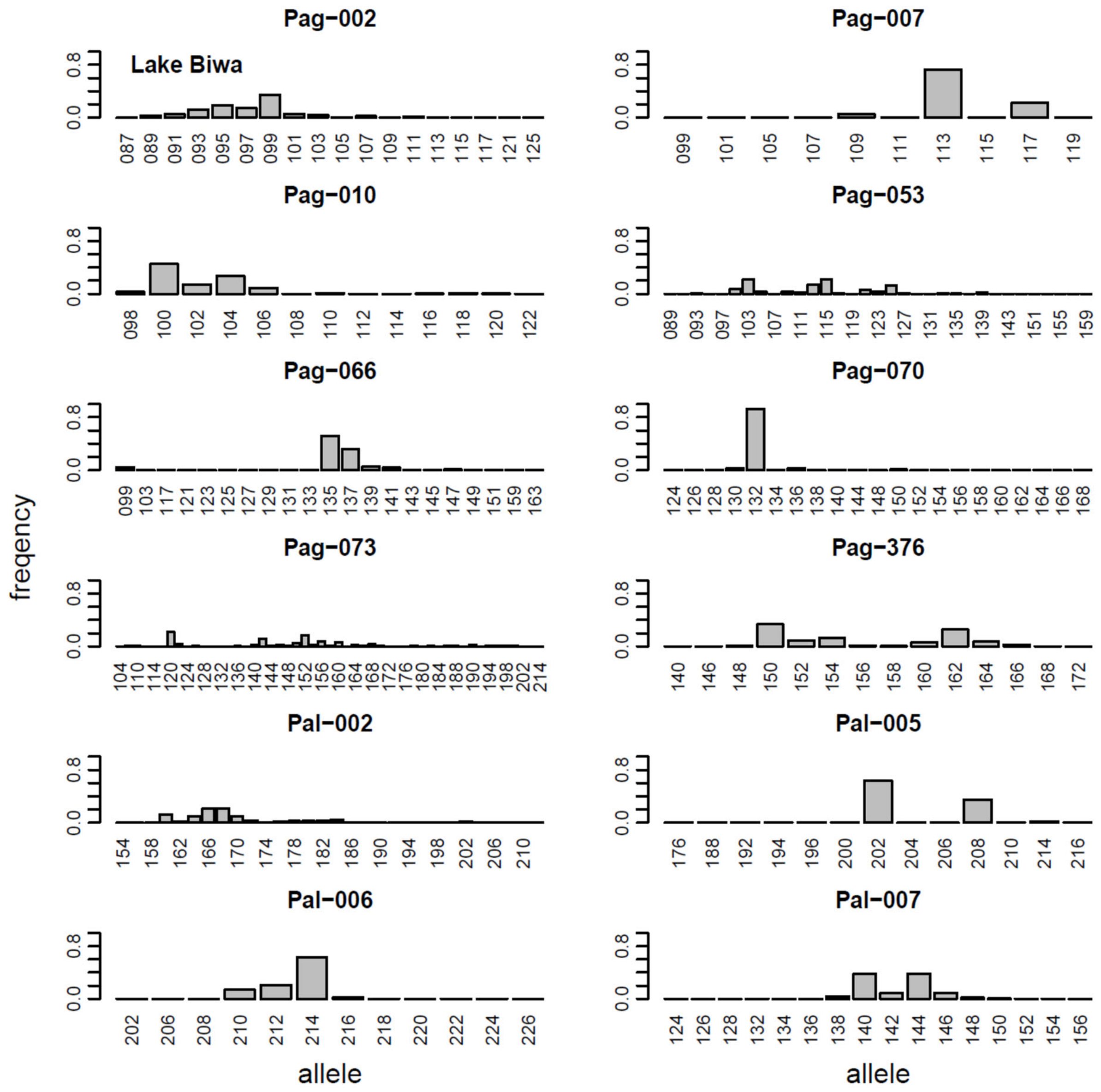


**Figure S4-2** Mean allele frequencies of landlocked form of Ayu at 12 microsatellite loci in Lake Biwa (20 populations, *n* = 718). Data are from Takeshima et al. (2016a)

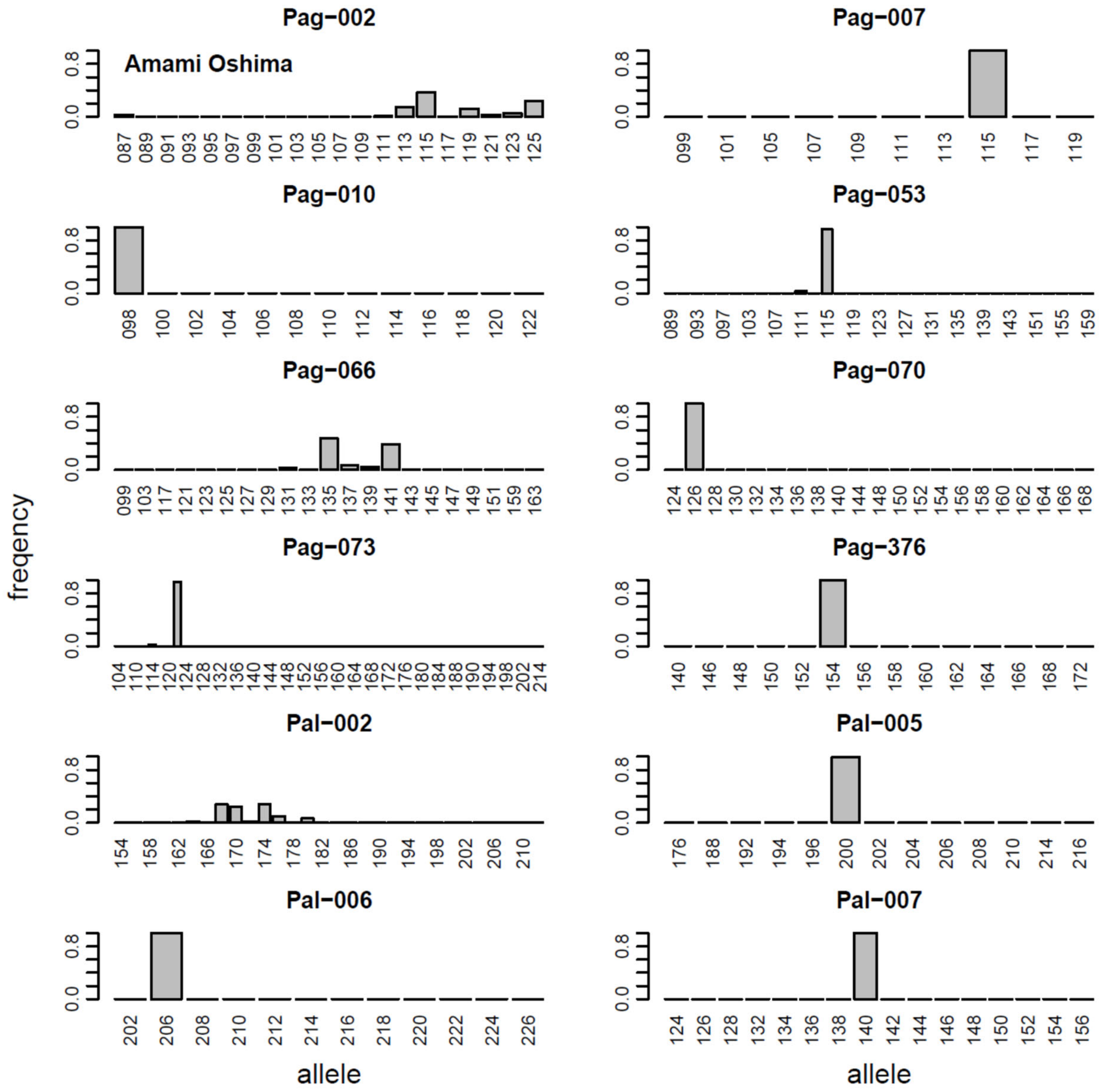


**Figure S4-3** Mean allele frequencies of Ryukyu Ayu at 12 microsatellite loci collected in Amami Oshima Island (2 populations, *n* = 63). Data are from Takeshima et al. (2016a)

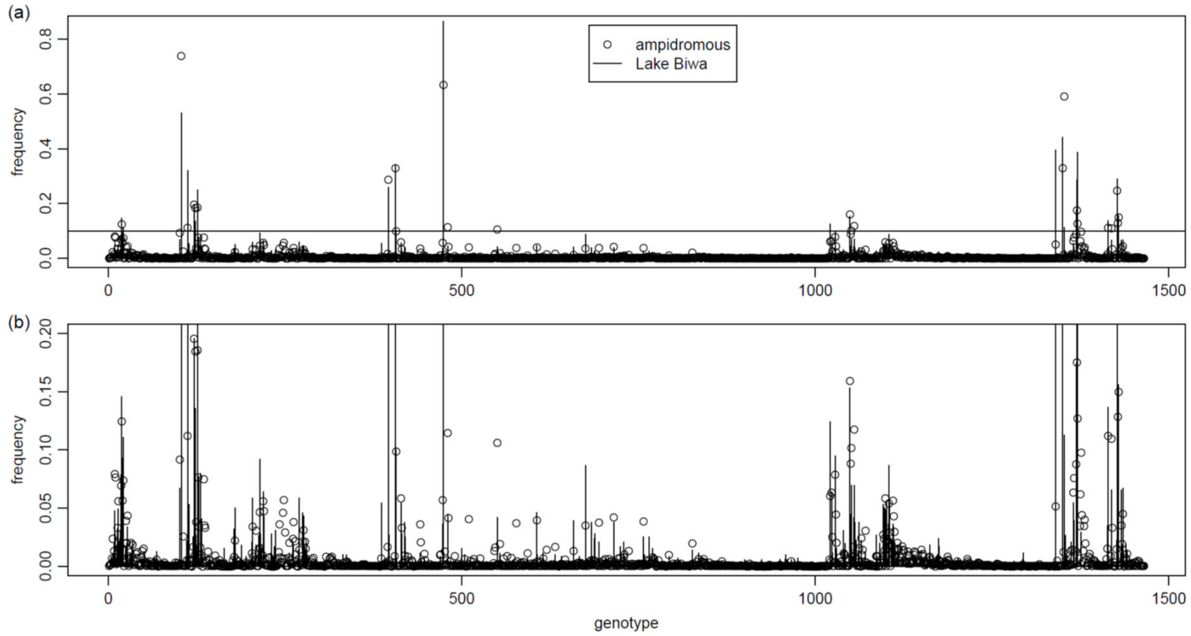


**Figure S5** Genotype frequencies for amphidromous and landlocked forms of Ayu in Japan. (a) Whole view. (b) Zoomed view for < 0.1 frequency. A total of 1465 genotypes were observed at 12 microsatellite markers in 98 amphidromous and 20 landlocked populations (*n* = 4746). Data are from Takeshima et al. (2016a)

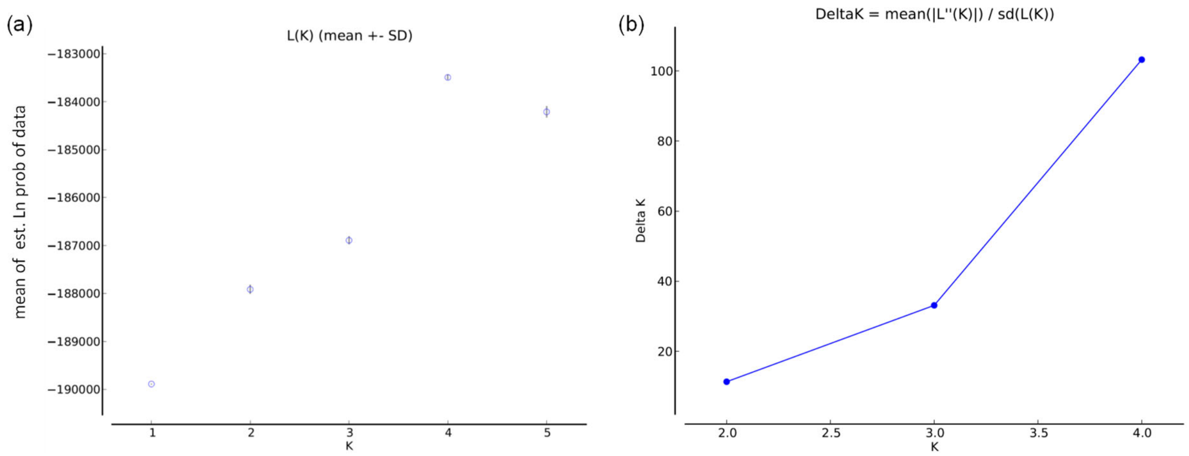


**Figure S6** Most probable number of clusters (*K*) as suggested by the method of Evanno (Evanno et al., 2005). (a) Ln Pr(data|*K*) and (b) *ΔK* were obtained using Structure Harvester (Earl & vonHoldt, 2012). Data are from Takeshima et al. (2016a)

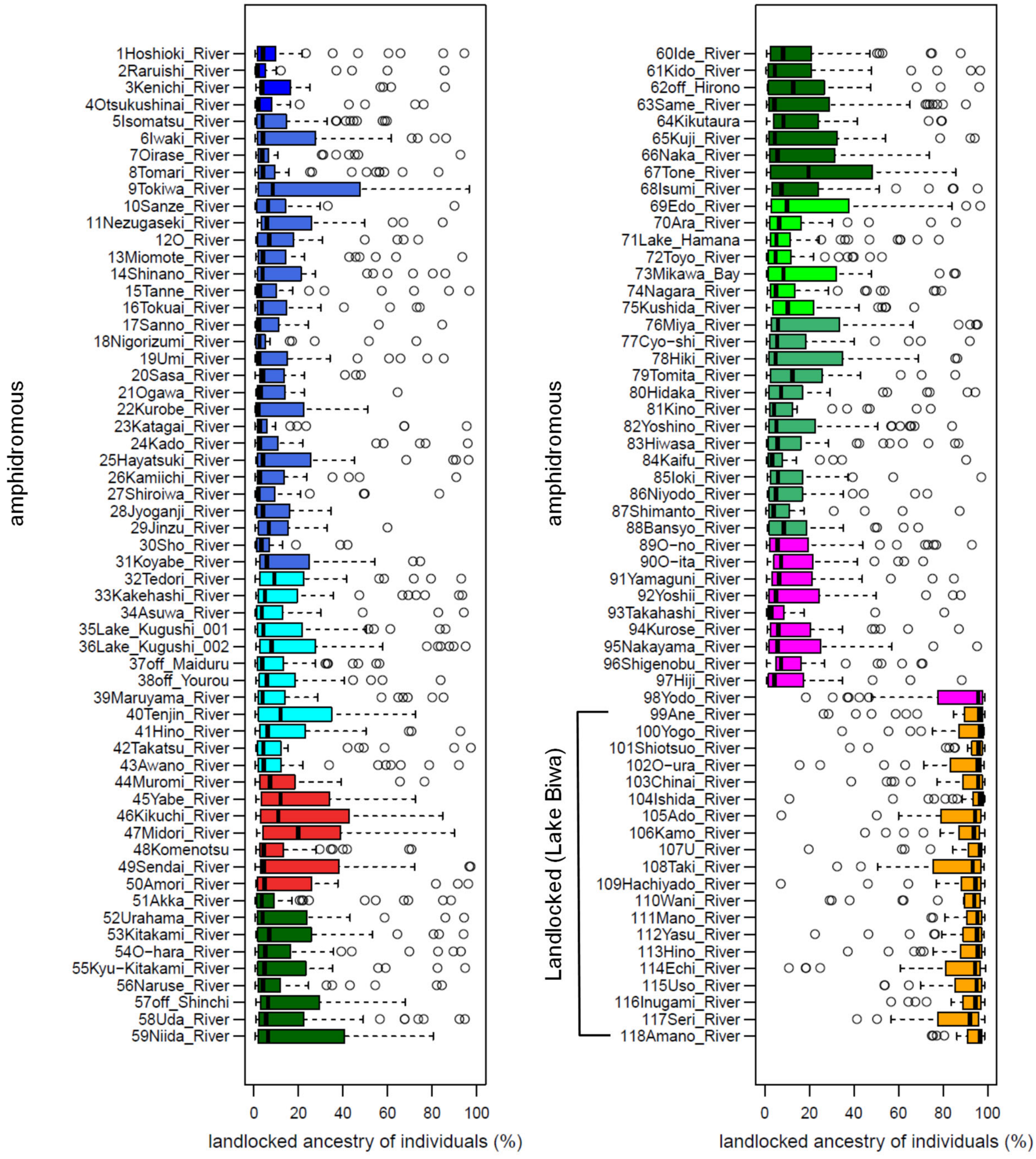


**Figure S7** Distribution of individual *q*-values of landlocked ancestry in 118 populations (shown in orange in Figure 5). Colours refer to sampling locations in Figure 2 (see Table S1 for sampling information). Data are from Takeshima et al. (2016a)

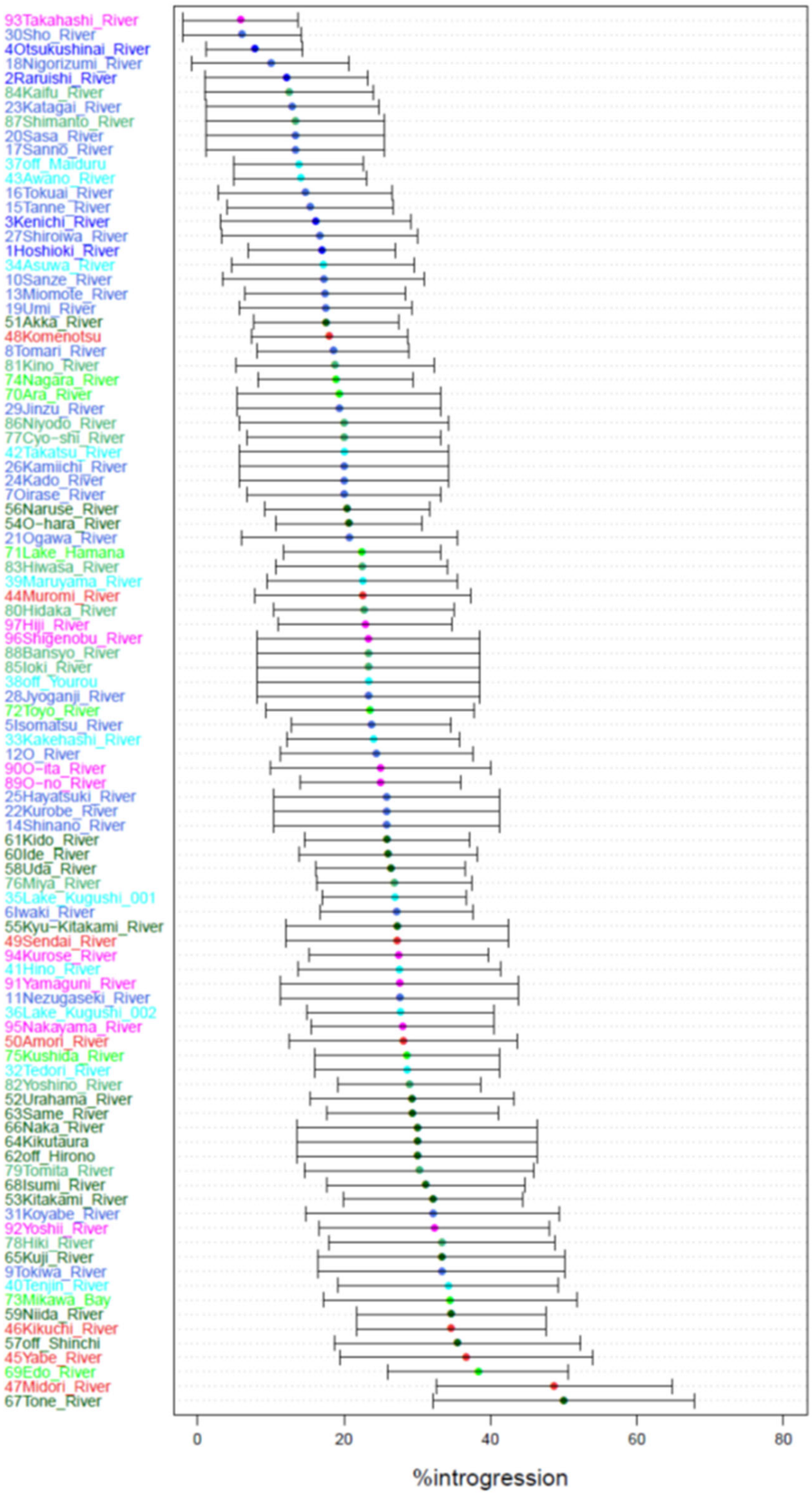


**Figure S8** Estimated introgression of landlocked Ayu ± 1.96× standard error in Japanese rivers. Colours refer to sampling locations in Figure 2 (see Table S1 for sampling information). Data are from Takeshima et al. (2016a)

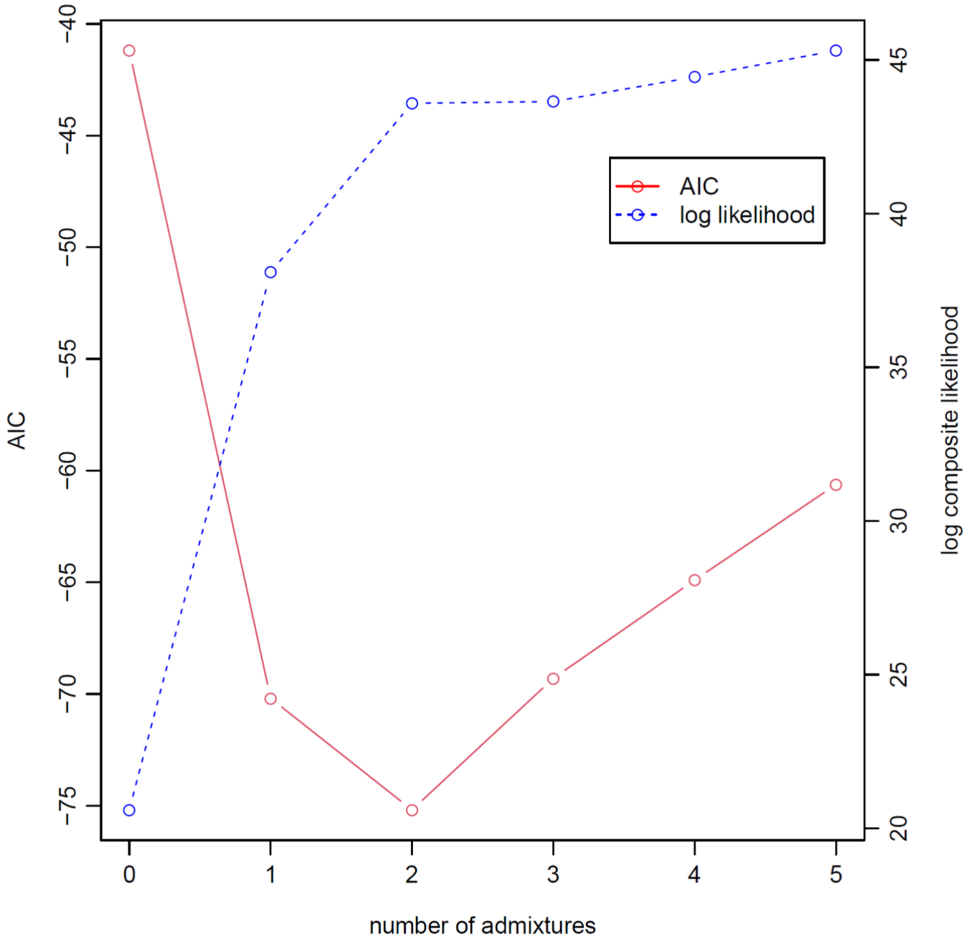


**Figure S9** Composite likelihood and Akaike Information Criterion (AIC) values for models of the number of admixture on the rooted tree from TreeMix analysis based on 12 microsatellite loci in 118 Ayu populations in Japan. Data are from Takeshima et al. (2016a)

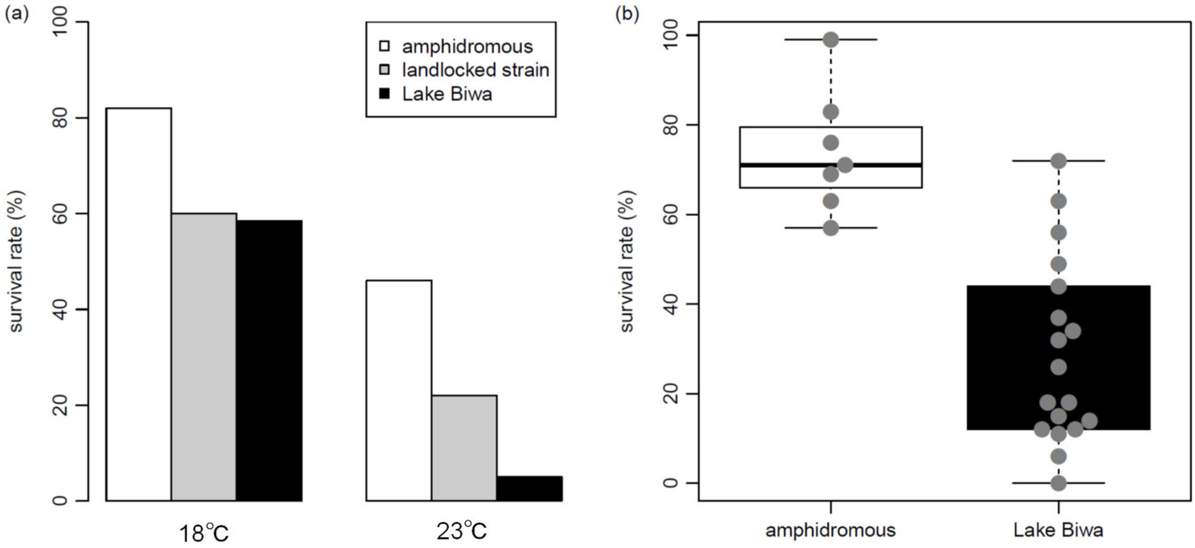


**Figure S10** Salinity tolerance experiments for Ayu larvae in seawaters. (a) Survival rates of larvae in seawater 20 days after hatching. The amphidromous form was born from wild fish that collected from Hiki River (Wakayama Pref.) and Kiso River (Aichi Pref.) in October and produced hatchlings of 6–7 mm total length (TL). The landlocked *strain* was born from wild fish that collected from Kiso River in September and produced hatchlings of 5 mm TL. Landlocked form larvae were born from wild parents collected in Lake Biwa. Drawn from Figure 1 in Tabata and Azuma (1986). (b) Survival rates of larvae in 80% artificial seawater (20°C) 24 hours after hatching. Salinity and temperature were set to values that drifting larvae experienced at a river mouth. Each grey point shows the survival rate of 30 individuals ($t = 6.28, df = 16.66, p = 9.05 \times 10^{-6}$, *t*-test). Drawn from Figure 3 in Iguchi and Yamaguchi (1994)
